# Identification of PAX6 and NFAT4 as the transcriptional regulators of lncRNA Mrhl in neuronal progenitors

**DOI:** 10.1101/2021.06.23.449546

**Authors:** Debosree Pal, Sangeeta Dutta, Dhanur P Iyer, Utsa Bhaduri, M.R.S Rao

## Abstract

LncRNA Mrhl has been shown to be involved in regulating meiotic commitment of mouse spermatogonial progenitors and coordinating differentiation events in mouse embryonic stem cells. Here we have characterized the interplay of Mrhl with lineage-specific transcription factors during mouse neuronal lineage development. Our results demonstrate that Mrhl is predominantly expressed in the neuronal progenitor populations in mouse embryonic brains and in retinoic acid derived radial-glia like neuronal progenitor cells. Mrhl levels are significantly down regulated in postnatal brains and in maturing neurons. In neuronal progenitors, a master transcription factor, PAX6, acts to regulate the expression of Mrhl through direct physical binding at a major site in the distal promoter, located at 2.9kb usptream of the TSS of Mrhl. Furthermore, NFAT4 occupies the Mrhl proximal promoter at two sites, at 437bp and 143bp upstream of the TSS. ChIP studies reveal that PAX6 and NFAT4 interact with each other, suggesting co-regulation of lncRNA Mrhl expression in neuronal progenitors. Our studies herewith are crucial towards understanding how lncRNAs are regulated by major lineage-specific TFs towards defining specific development and differentiation events.

**Summary statement:** Transcriptional regulation of lncRNA Mrhl by multiple lineage-specific transcription factors in neuronal progenitors highlights context-dependent regulation important for lineage specification.

## Introduction

Non-coding RNA dominates the eukaryotic transcriptome contributing up to 80-90% of it (Zhang *et. al*., 2019 a). Long non-coding RNAs (lncRNA) are classified as a major class of non-coding RNAs with a length greater than 200 nucleotides, without any or little translational activity (Wilusz *et. al*. 2009; Jarroux *et. al*., 2017). LncRNAs which do show little translational activity has been reported to possess small open reading frame which encodes for small or micro peptides of fundamental biological importance (Hartford & Lal, 2020; Xing *et. al*., 2020). Although most of the long non-coding RNAs (lncRNAs) are not well conserved, their highly specific spatial-temporal expression has highlighted their complex roles in diverse biological functions during development and differentiation. And their dysregulation has been associated with several diseases including cancer and neurological disorders (Iyer *et. al*., 2015; Kopp & Mendell, 2018; Marchese *et. al*., 2017; Wapinski & Chang, 2011; Sun & Kraus, 2015; Li *et. al*., 2019). Several studies have verified the importance of long non-coding RNAs as functional regulators. To regulate their targets LncRNAs are known to use a wide variety of mechanisms both at transcriptional and post transcriptional levels (Chen & Tergaonkar, 2020; Rosa & Ballarino, 2016; Yoon *et. al*., 2013; Akhade *et. al*., 2017).

During development of a mammalian system, a synchronous expression and repression of genes leads to a staggering variety of cells with various functional activities. This remarkable orchestration of genetic component is most evident in an extremely complex system of mammalian neural development and brain with its intricate cellular architecture, maintenance and function. However, regulators behind this fine coordination are still elusive. With a large number of highly specific lncRNAs expressed in the brain, their contribution in neuronal development is undeniable (Zhao *et. al*., 2020; Roberts *et. al*., 2014; Briggs *et. al*., 2015; Hart & Goff, 2016; Mercer *et. al*., 2008; Derrien *et. al*., 2012; Kadakkuzha *et. al*., 2015). Data from GENCODE v7 catalog suggest 40% of most differentially expressed lncRNAs are specific for brain (Derrien *et. al*., 2012). A notable aspect is their distinct neuroanatomical loci along with developmental and stage specific expression as evident in many studies (Liau *et. al*., 2021; Lv *et. al*., 2013; Antoniou *et. al*., 2014; Dinger *et. al*., 2008; Mohamed *et. al*., 2010; Hezroni *et. al*., 2020). For example, in a study by Goff. *et. al*., they used mutant mouse models for 13 lncRNAs (Lincenc1, Eldr, Pantr1, Pantr2, Tug1, Peril etc.) and assessed their expression pattern from E14.5 to adult brain revealing high variability in expression pattern for each lncRNA both spatially and temporally (Goff *et. al*., 2015). Along with characteristic lncRNA expression signatures in sub-regions of the brain and specific neuronal populations, studies have also shown lncRNA-mRNA pairs, sharing a genomic locus, to be displaying specific coexpression in a region-specific manner. For example, lncRNA GM9968 and DNA binding protein gene *Zbtb20* show high enrichment in hippocampus (Kadakkuzha *et. al*., 2015). Transcriptome analysis by Lv. *et. al*. has revealed the development stage specificity of lncRNAs in mouse (Lv *et. al*., 2013). They have annotated large number of lncRNAs to the embryonic brain. Interestingly they found that expression of most them were very low or non-evident in the adult brain, suggesting their significance during embryonic brain development (Lv *et. al*., 2013).

LncRNAs have been shown to undergo regulated expression during development and differentiation (Fullard *et. al*., 2018). Studies have verified dynamic expression and regulation of many lncRNAs in conjunction with stage-specific transcription factors (TFs) during *in-vitro* differentiation scenarios like of mouse embryonic stem (ES) cells to embryoid bodies (EB) or during retinoic acid mediated neuronal differentiation of ES cells (Antoniou *et. al*., 2014; Dinger *et. al*., 2008). Observations from these studies indicate association of pluripotent factors like SOX2, OCT4 and NANOG at the promoter of lncRNAs, modulating their expression during development. For example, lncRNA AK028326 is activated by OCT4 while lncRNA AK141205 is repressed by NANOG resulting in the modulation of mouse ES cell pluripotency (Mohamed *et. al*., 2010). This tightly controlled differential expression pattern of lncRNAs is thus important for the delicate balance between lineage commitment and maintenance of pluripotent status. Results from loss and gain of function studies involving lncRNAs also show the vice versa to be true. Knockdown and over expression of lncRNAs alters the expression of transcription factors regulating them, leading to disruption of lineage specification. For example, using inducible knockdown of lncRNAs Rik-201 and Rik-203 in mouse ES cells, Zhang *et. al*. proved their importance in neural differentiation. Repression of lncRNAs Rik-201 and Rik-203 contributes to inhibition of neural differentiation by modulating the expression of SOX6 (Zhang *et. al*., 2019 b). A separate study identifies lncRNA Sox2OT as a promoting factor for cortical neural progenitors by repressing the expression of pluripotent gene *Sox2* via physical interaction with multifunctional transcriptional regulator YY1 (Knauss *et. al*., 2018). Another example of regulation of *Sox2* expression by a lncRNA which alters the neural stem fate is by lncRNA Rmst which physically interacts with SOX2 to alter the outcome (Ng *et. al*., 2013).

LncRNA Mrhl (meiotic recombination hotspot locus), a lncRNA that has been studied extensively by our group, is the focus of our present study (Nishant *et. al*., 2004; Ganesan & Rao, 2008; Arun *et. al*., 2012; Akhade *et. al*., 2014; 2016; 2017; Pal *et. al*., 2021). Mrhl is an intronic and single exonic lncRNA with a length of 2.4 kb. This nuclear restricted lncRNA is transcribed from the 15^th^ intron of *Phkb* gene in mouse and shows syntenic conservation in humans (Ganesan & Rao, 2008; Fatima *et. al*., 2019; Choudhury *et. al*., 2020, Preprint). LncRNA Mrhl is found to be a negative regulator of WNT signaling via its interaction with p68/DDX5 RNA helicase in mouse spermatogonial cells (Ganesan & Rao, 2008; Arun *et. al*., 2012; Akhade *et. al*., 2016). LncRNA Mrhl has also been shown to regulate *Sox8* at chromatin level signifying its role in meiotic commitment of mouse spermatogonial progenitor cells (Akhade *et. al*., 2014). Our latest finding from transcriptomics and genome-wide occupancy studies of Mrhl in mouse ES cells has revealed its role as a chromatin regulator of cellular differentiation and development genes (Pal *et. al*., 2021).

Taking clue from our recent study (Pal *et. al*., 2021) where a large number of genes related to neural development and differentiation were dysregulated upon Mrhl downregulation, we explored the dynamic expression pattern of lncRNA Mrhl and its transcriptional regulation during mouse neuronal lineage development in the present study. We have shown that Mrhl is predominantly expressed in the neuronal progenitor populations in the ventricular-subventricular zones of developing mouse embryonic brains. We have adopted an *in vitro* model system of retinoic acid mediated derivation of radial-glia like neuronal progenitor cells (NPCs) from mouse ES cells. Mrhl shows up regulation of expression in these NPCs. Levels of lncRNA Mrhl is significantly down regulated in postnatal brains and in maturing neurons suggesting its significance in embryonic neuronal lineage development. We have also identified PAX6, a master transcription factor in NPCs, to be a regulator of Mrhl transcription. Further exploration of mechanism reveals that PAX6 can do so by physically binding to the promoter of Mrhl at multiple sites of action. A major site of action for PAX6 on Mrhl promoter is situated 2.9kb upstream of the TSS where it binds directly and regulates Mrhl in NPCs. Three other minor sites are present at 1.27kb, 672bp and 622bp upstream of the TSS of Mrhl wherein regulation is proposed to be by PAX6 and/or its isoform PAX6(5A) and potentially in conjunction with other co-factors. NFAT4, which is another important transcription factor in neuronal development, was also found to be enriched at the promoter region of Mrhl at two sites in the proximal promoter, 437bp and 143bp upstream of the TSS. ChIP studies verifies interactions of PAX6 and NFAT4 with each other leading to the assumption that PAX6 might regulate Mrhl in coordination with NFAT4 in NPCs. This study thus reveals regulated expression of lncRNA Mrhl by multiple and specific transcription factors during neuronal lineage development.

## Results

### Mrhl is predominantly expressed in embryonic stages of the mouse brain and in neuronal progenitor population

Our recent study on the transcriptome analysis of mouse ES cells following down regulation of Mrhl lncRNA showed an over representation of neuronal lineage-specific genes and networks to be dysregulated (Pal *et. al*., 2021). Based on this observation we initiated this study towards addressing the role of Mrhl in neuronal lineage specification. To begin with, we analyzed its expression level in the mouse brain tissues of both embryonic and postnatal mice. We performed qRT-PCR on brains from embryonic stages E10.5-E18.5 and postnatal stages P0-P40 and observed that Mrhl expression is predominantly abundant in stages E12.5-E18.5 (**Fig. 1A**). Furthermore, Mrhl showed abrupt decrease in expression level from postnatal stage P0 onwards. At around E8 of brain development, neuroepithelial cells of the neural tube give rise to radial glia cells (RGCs), the future NPCs of the brain. RGCs undergo rapid proliferation and specification into basal progenitors which in turn give rise to neurons, with a peak of neurogenesis at E14 (Yao & Jin, 2014). Since we observed the first peak of Mrhl expression at E14.5 in our studies, we focused to understand its role at this stage. We isolated brain tissue from E14.5 embryos and divided them into fore-, mid- and hind-brain regions based on their gross anatomy. We confirmed the purity of the three regions obtained based on forebrain specific marker FoxG1, midbrain specific marker En1 and hindbrain specific marker Gbx2 (Kirkeby *et. al*., 2012; Su *et. al*., 2018). Pax6 was used a control marker owing to its defined expression pattern across the three regions of the brain. We observed that Mrhl is expressed ubiquitously across all the three regions of the mouse brain at E14.5 (**Fig. 1B**). Whilst Mrhl may have specific roles to play across all regions of the developing brain, we concentrated on the forebrain development. To further narrow down into the specific cell types in which Mrhl might be involved in the forebrain, we performed RNA FISH for Mrhl along with IF for PAX6 as the marker, on whole forebrain sections at E14.5. We found that Mrhl is expressed specifically in the ventricular/subventricular zone of the brain at this stage (**Fig. 1C**). This was further confirmed by analyzing publicly available ChIP-Seq datasets [GSE93009] for RNA Pol II, RNA Pol II-Ser5p (initiation polymerase), transcription activation mark, H3K4me3 and transcription repression mark, H3K27me3, on the proximal 1Kb promoter of Mrhl in Nestin+ NPCs isolated from E15.5 cortices (Liu *et. al*., 2017). The analysis showed us distinct enrichments for RNA Pol II, initiating RNA Pol II and H3K4me3 near the TSS for Mrhl (**Fig. 1D**). Thus, we concluded that Mrhl is predominantly expressed in the neuronal progenitor population of the developing embryonic mouse brain.

**Figure 1.**
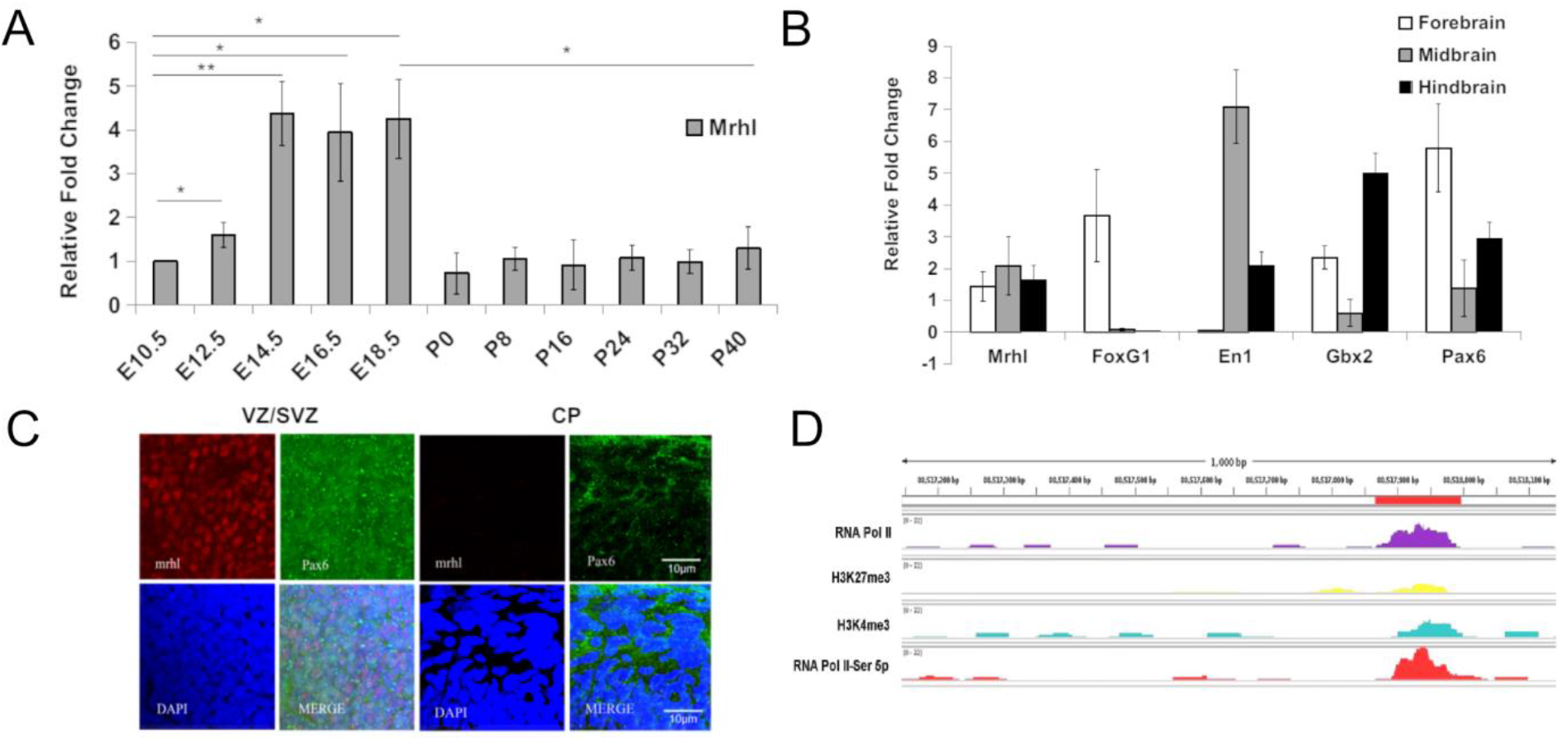
Mrhl is predominantly expressed in the neuronal progenitor population of the developing mouse embryonic brain. (A) qPCR data for Mrhl expression in embryonic and postnatal mouse brains. (B) qPCR for Mrhl expression across different regions of E14.5 brain along with region-specific markers. (C) RNA FISH for Mrhl and IF for PAX6 in E14.5 brain sections shows predominant expression of Mrhl in ventricular-subventricular (VZ/SVZ) zones of the brain concomitant with PAX6 nuclear expression. CP = cortical plate. (B) Analysis of available ChIP-Seq datasets for enrichment of RNA Pol II or histone modifications in Nestin+ cells from E15.5 cortices. Error bars indicate standard deviation from three independent experiments, with replicates in each (n=6-20). *p<0.05, **p<0.01, ***p<0.001, student’s t-test; Scale bar = 10 μm.

### Mrhl expression pattern in mouse brain is recapitulated in an *in-vitro* differentiation model

We adopted the well-established RA induced *in vitro* differentiation model system to further elucidate Mrhl as a molecular player in NPCs. We generated EBs from mouse ES cells and treated them with retinoic acid (RA). This method has been shown to generate PAX6+ RGC-like NPCs that possess the capability to generate neurons *in vivo*, as shown by the study of Bibel *et. al*. and Plachta *et. al*. (Bibel *et. al*., 2007; Plachta *et. al*., 2004). We observed that Mrhl is up regulated in expression in RA treated EBs as compared to vehicle treated EBs (**Fig. 2A**). An analysis for various NPC markers such as Pax6 and Nestin as well as neuronal markers such Tuj1 and Ascl1 confirmed our *in vitro* neuronal lineage differentiation system (**Fig. S1A, B**). We next differentiated these *in vitro* derived neuronal progenitors into neurons and observed that Mrhl expression is abruptly down regulated from 12 hours post neuron formation (**Fig. 2B**). We further confirmed our observations through RNA FISH for Mrhl and IF for NESTIN on cryosections of RA versus vehicle treated EBs. RA treated EB sections showed noticeable expression for Mrhl as compared to vehicle treated sections (**Fig. 2C**). Similarly, Mrhl showed significant reduction in expression from early neuron stage at 12 hours (**Fig. 2D, S1C**). We also performed ChIP-qPCR on RA treated EBs and neurons at 24 hours of differentiation for H3K4me3, an active transcription mark, and the results indicated a decreased association of this histone mark at the Mrhl promoter in maturing neurons (**Fig. 2E**). Our observations thus confirm that Mrhl is preferentially expressed in NPCs.

**Figure 2.**
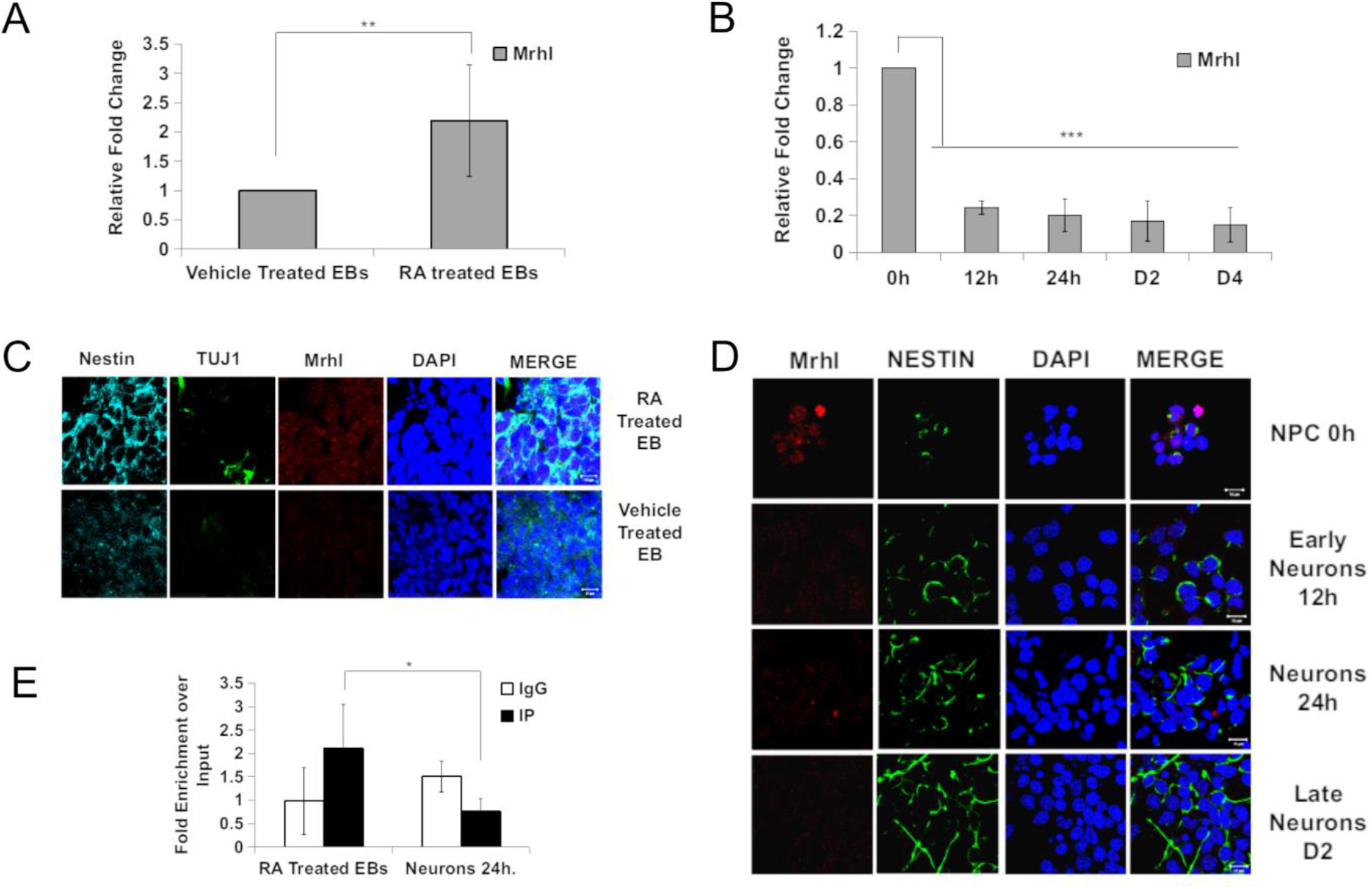
Mrhl expression in embryonic brain is recapitulated in an *in vitro* model. (A, B) qPCR analysis of Mrhl expression in RA versus vehicle treated EBs and 12 hours to D4 old maturing neurons, respectively. (C, D) RNA FISH for Mrhl and IF for NPC marker NESTIN and maturing neuron marker TUJ1 in EB sections and neurons. (E) ChIP-qPCR for H3K4me3 on proximal Mrhl promoter in RA treated EBs versus neurons at 24 hours. Error bars indicate standard deviation from three independent experiments, with replicates in each (n=6). *p<0.05, **p<0.01, ***p<0.001, student’s t-test; Scale bar = 10 μm.

### A master transcription factor PAX6 was identified as a regulator of Mrhl in NPCs

Next, we were interested in understanding how the transcriptional activity of Mrhl is regulated in neuronal progenitors. LncRNAs act in close coordination with TFs to not only regulate target genes but also be regulated themselves. LncRNA Rmst, also known as Ncrms, has been shown to be positively regulated by PAX2, a TF involved in midbrain and cerebellum development (Bouchard *et. al*., 2005) whereas it is negatively regulated by REST, during neurogenesis (Ng *et. al*., 2013), emphasizing the context-dependent transcriptional regulation of lncRNAs by major TFs. STAT3 has been shown to up regulate the expression of lncRNA Hoxd-As1 at the transcription level, contributing to invasion and metastasis in hepatocellular carcinoma (Wang *et. al*., 2017). During male germ cell meiotic commitment, Mrhl is repressed in transcription by the WNT signaling pathway effector, TCF4/β-CATENIN complex along with the corepressor CTBP1, via interaction at its promoter (Akhade *et. al*., 2016).

We used a combination of GPMiner (Lee *et. al*., 2012) and JASPAR (Mathelier *et. al*., 2014) programs to gain insights into the probable transcriptional regulators of Mrhl in neuronal progenitors. An analysis for 3kb upstream of TSS region for Mrhl revealed predicted binding sites for diverse lineage specific TFs **(Fig. S2A)**. With respect to the neuronal lineage, TCF4 (172bp), PAX6 (2904bp, 1276bp, 672bp and 622bp), RBPJ-K (484bp), NFAT (437bp and 143bp) and MEIS1 (477bp) were of noticeable importance (**Fig. S2A and Supplementary File 1**). Initially, we focused on TCF4, an effector of the WNT pathway owing to our earlier studies on WNT mediated regulation of Mrhl in B-type spermatogonial progenitors. Additionally, the WNT pathway has been shown to be involved in context-dependent roles in neural development (Chenn & Walsh, 2002; Machon *et. al*., 2003; Woodhead *et. al*., 2006). Towards this end, we performed IF for β-CATENIN on RA treated versus vehicle treated EBs as well as during neuronal differentiation over a time period of 2h-24h **(Fig. S2B)**. We observed no translocation of β-CATENIN from membrane to the nucleus, a hallmark of non-activation of the WNT pathway. Hence, we ruled out TCF4 as a potential transcriptional regulator for Mrhl in RA derived NPCs. Our studies also suggest that WNT signaling may not have a role to play in RA derived NPCs. We additionally probed into RBPJ-K, an effector of the NOTCH pathway, since NOTCH signaling has been extensively implicated in the proliferation and maintenance of radial glia progenitors in the brain (Gaiano *et. al*., 2000; Mizutani *et. al*., 2007; Imayoshi *et. al*., 2010; Sakamoto *et. al*., 2003). We performed IF for NICD in vehicle treated versus RA treated EBs and observed translocation of NICD to the nucleus of RA derived NPCs **(Fig. S2C)**, indicating activation of the NOTCH pathway. However, subjecting RA treated EBs to the NOTCH pathway inhibitor DAPT, showed a reduction in expression for the known target Hes5 but not for Mrhl **(Fig. S2D)**, thereby ruling out RBPJ-K as the potential regulator for Mrhl. We also did not see changes in expression levels for other neuronal markers such as Pax6 and Tuj1, suggesting in RA derived NPCs, NOTCH signaling maybe acting through other pathways altogether to regulate progenitor physiology. We then focused on PAX6 whose role in neural development w.r.t NPCs and neurogenesis has been well established (Sansom *et. al*., 2009; Thakurela *et. al*., 2016). We collected ChIP-Seq datasets available for PAX6 in E12.5 mouse embryonic forebrain [GSE 66961] (Sun *et. al*., 2015) and combined them with a FIMO analysis for PAX6 motifs (Xie & Cvekl, 2011; Xie *et. al*., 2013). We narrowed down four binding sites on the promoter of Mrhl as candidates for PAX6 mediated regulation (**Fig. 3A**, sites 1, 2, 3 and 4). The FIMO analysis further revealed the presence of motifs 1-1 and 2-1 in site 1, motifs 1-3 and 3-3 in site 2 and motif 4-1 in sites 3 and 4 (**Fig. 3B-D, Table 1, Supplementary File 1**). ChIP-qPCR for PAX6 corresponding to sites 1 to 4 on Mrhl promoter showed that PAX6 occupies all sites in E14.5 brains and sites 1, 3 and 4 in RA treated EBs (**Fig. 3E-F, S3A**). We concluded here that PAX6 is potentially regulating Mrhl at the transcriptional level in NPCs.

**Figure 3.**
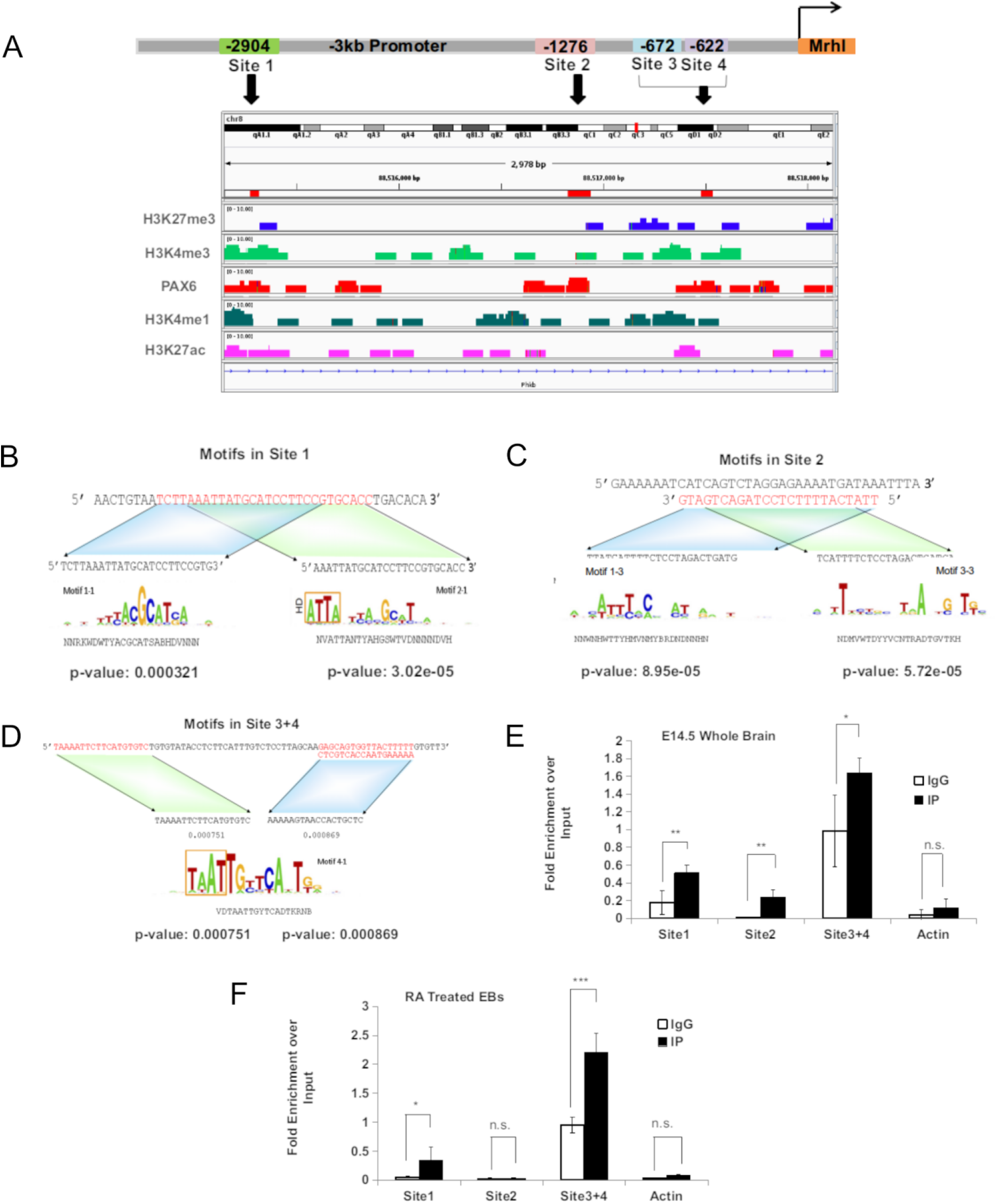
Identification of potential transcriptional regulators for Mrhl during neuronal lineage development. (A) ChIP-Seq analysis for PAX6 and histone modifications on 3kb promoter of Mrhl, red bars and black arrows show our sites of interest. (B-D) Motifs for PAX6 on Mrhl promoter sites 1 to 4 as revealed by FIMO analysis. (E, F) ChIP-qPCR analysis for PAX6 enrichment on Mrhl promoter in E14.5 whole brain and RA treated EBs. Error bars indicate standard deviation from three independent experiments, with replicates in each (n=6). *p<0.05, **p<0.01, ***p<0.001, student’s t-test.

**Table 1:**
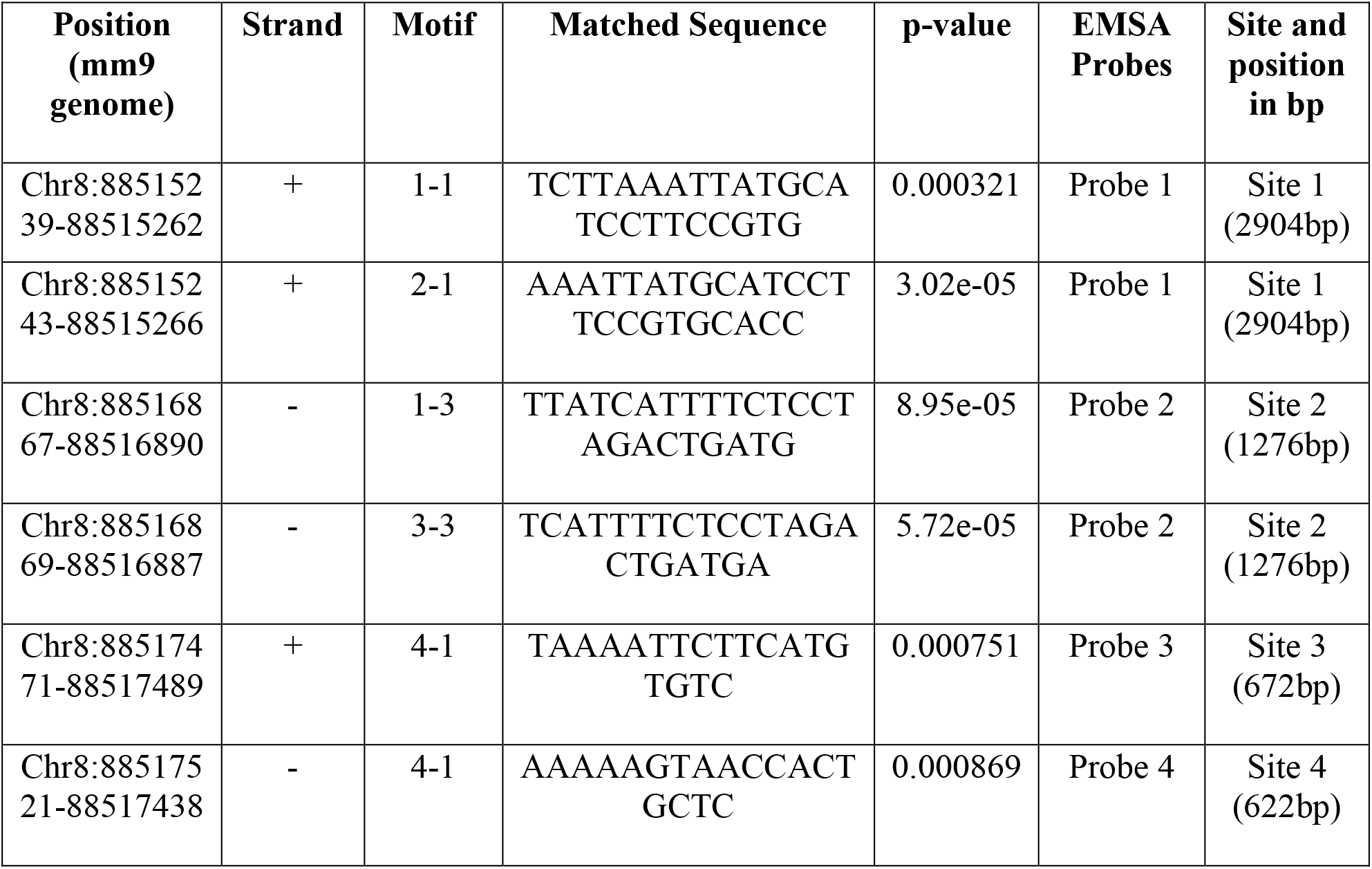
PAX6 binding sites and corresponding EMSA probes used for Mrhl promoter, 3kb upstream from TSS.

### PAX6 physically binds to the distal promoter and regulates Mrhl in NPCs

We next performed an in-depth analysis involving EMSA, luciferase and knockdown assays to understand the regulation of Mrhl by PAX6 at these four sites. We expressed and purified fulllength PAX6 and its isoform PAX6(5A) (**Fig. S3B**) for our EMSA experiments (**Table 1**). We observed that PAX6 binds directly to site 1 on Mrhl promoter strongly, to site 2 weakly and does not bind to sites 3 and 4 (**Fig. 4A**). Elaborate studies by Xie and Cvekl (Xie & Cvekl, 2011) have shown that PAX6 binds to nine novel motifs, only one of which corresponds to the consensus. Furthermore, PAX6 and PAX6(5A) bind to these motifs with varying affinities depending on the biological context. Since, we observed occupancy of PAX6 in our ChIP results on sites 2, 3 and 4 and since it has already been demonstrated that motif 1-3 in site 2 and motif 4-1 in sites 3 and 4 are bound by PAX6(5A) with moderately high affinity (Xie & Cvekl, 2011), we hypothesized, sites 2, 3 and 4 might be bound by the isoform PAX6(5A). PAX6(5A) differs from PAX6 in that it has a 14 amino acid insertion in the PAI domain, thereby its differential binding to the various motifs. Furthermore, studies have shown that the ratio of PAX6:PAX6(5A) shifts from 8:1 to 3:1 from stages E12.5 to E14.5 of development (Pinson *et. al*., 2005). However, our EMSA studies showed no binding for PAX6(5A) to any of these sites (**Fig. S3C**). We then carried out luciferase assays in P19EC cells which lack expression of endogenous PAX6 using various constructs of Mrhl promoter harboring one or the other sites as shown in **Fig. 4B-D** in the presence of ectopically expressed PAX6 or PAX6(5A). Sites 3 and 4 did not elicit any luciferase activity. However, both sites 2 and 1 displayed luciferase activity in the presence of both the isoforms. Furthermore, luciferase assays in the presence of both the isoforms revealed highest activity at a ratio of 4:1 for PAX6:PAX6(5A) for the Mrhl promoter (**Fig. 4E**). Finally, we generated doxycycline inducible PAX6 knockdown mouse ES cell lines and induced the knockdown of PAX6 at day 5 of RA mediated differentiation of EBs and scored for Mrhl expression. We observed a reduction in Mrhl expression concomitant with reduction of PAX6 expression (**Fig. 4F**). From these studies we infer that PAX6 transcriptionally regulates Mrhl in neuronal progenitors through direct physical binding at site 1 on the distal promoter of Mrhl. Site 2 may also be a point of regulation by PAX6/PAX6(5A) as is evidenced by our ChIP and luciferase assays probably in conjunction with other TFs/cofactors. It has been shown that other TFs such as SOX2 act in conjunction with PAX6 to regulate enhancers of target genes in lens (Kamachi *et. al*., 2001). Sites 3 and 4 showed enrichment for PAX6 in our ChIP assays but no binding or activity in our EMSA and luciferase assays, respectively, suggesting regulation via these two sites might be highly context dependent.

**Figure 4.**
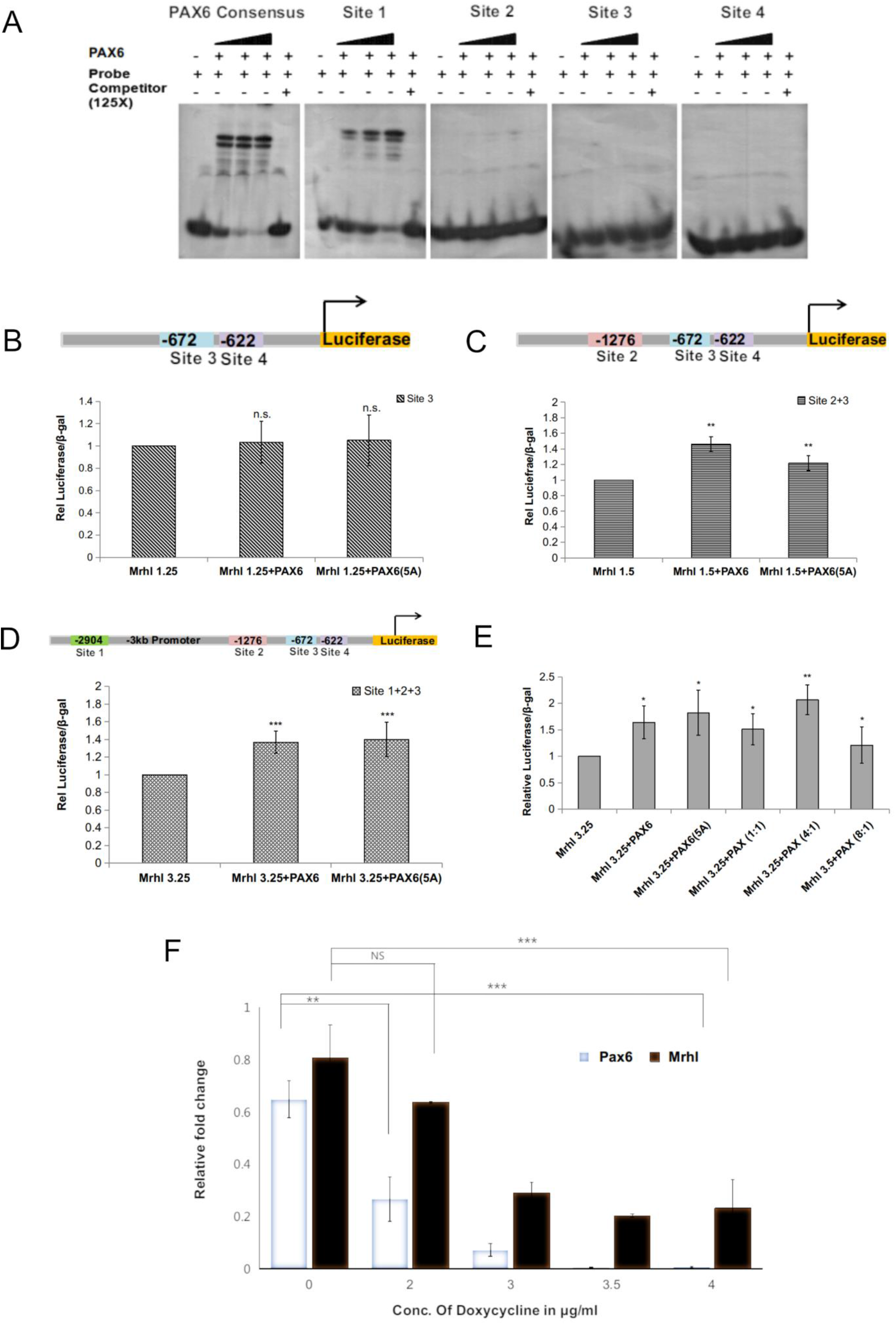
PAX6 is a major transcriptional regulator of Mrhl in neuronal progenitors. (A) EMSA for studying direct interaction of PAX6 on the Mrhl promoter sites 1-4. (B-E) Luciferase assays for different constructs of Mrhl promoter in the presence of exogenously expressed PAX6 and PAX6(5A). (F) Knockdown assay showing decrease in Mrhl expression upon inducing knockdown of PAX6. Error bars indicate standard deviation from three independent experiments, with replicates in each (n=6). *p<0.05, **p<0.01, ***p<0.001, student’s t-test.

### Regulation of Mrhl by NFAT4 in NPCs at the proximal promoter

The experiments described above clearly demonstrate that PAX6 regulates Mrhl expression in NPCs, majorly through the upstream distal site 1 at 2.9kb in the promoter of Mrhl. We argued that there must be other TFs that act to regulate Mrhl via its proximal promoter and they may act in conjunction with PAX6. From our earlier analysis of the 3kb upstream of TSS region for Mrhl, we observed that NFAT and MEIS 1 have potential binding sites in the proximal promoter of Mrhl. MEIS1 has been implicated in innervation in the neural crest (Bouilloux *et. al*., 2016), during RA mediated neural differentiation of mouse P19EC cells (Yamada *et. al*., 2013) and in granule cells during development of the cerebellum (Owa *et. al*., 2018). NFAT has been implicated in being involved in various aspects of neural development such as axon growth, neuronal survival and synapse communication (Vihma *et. al*., 2016; Wild *et. al*., 2019). NFAT showed two possible binding sites at 143bp (BS 1) and 437bp (BS 2) upstream from Mrhl TSS, while MEIS1 binding site was found at 477bp upstream. Since NFAT has four isoforms, all of which are involved in neural pathways, we first established which isoform of NFAT could be regulating Mrhl in NPCs through PCR studies. We observed that only NFAT4 is expressed in both E14.5 embryonic brain as well as in EBs (**Figure 5A and S4A**). We further observed that NFAT4 exhibits the highest expression amongst all isoforms in embryonic brains (expression was compared w.r.t P4 brain in each case, **Figure S4B**). We next performed ChIP-PCR to verify the enrichment of both NFAT4 and MEIS1 at the promoter site of Mrhl. Our result show highly significant enrichment of NFAT4 at both the predicted binding sites (**Figure 5B**). Although MEIS 1 also showed significant enrichment at the Mrhl promoter, the enrichment was ~ 4-fold less as compared to NFAT4. Therefore, we focused on NFAT4 herewith and tried to determine if it interacts with PAX6 for a possible co-regulation of Mrhl. Co-ChIP western blots clearly demonstrated enrichment of NFAT4 during PAX6-IP (**Figure 5C**) and vice-versa (**Figure 5D**). Findings from these results lead us to the conclusion that PAX6 regulates Mrhl at the distal promoter whereas NFAT4 acts to possibly regulate Mrhl at the proximal promoter in NPCs. There is a further possibility that PAX6 and NFAT4 act to co-regulate lncRNA Mrhl in NPCs towards regulating cell fate decisions (**Figure 5E-F**).

**Figure 5.**
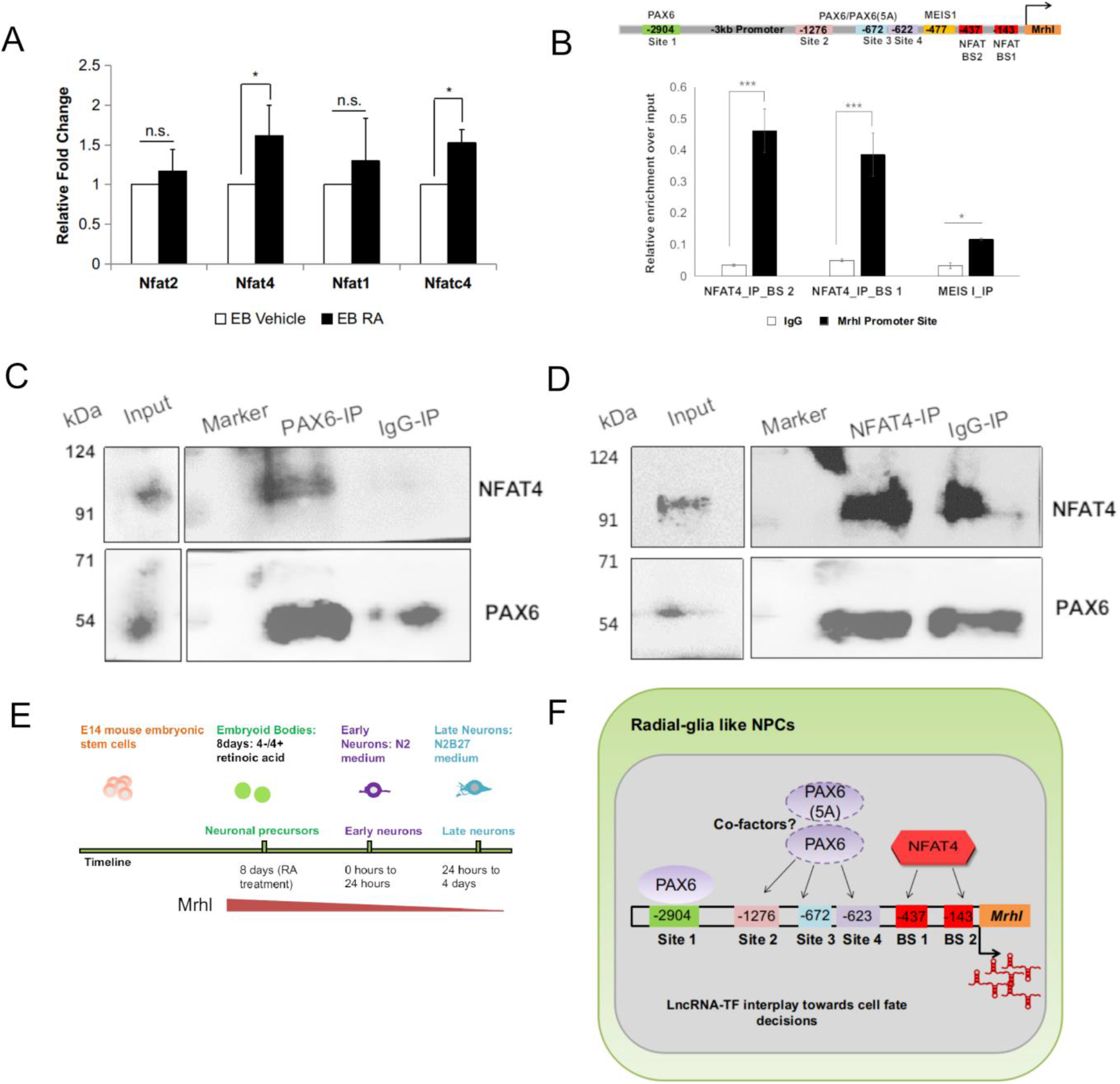
(A) Expression pattern of various NFAT isoforms vehicle versus RA treated EBs. (B) ChIP-qPCR for NFAT4 and MEIS1 binding on Mrhl promoter. (C) ChIP-WB for NFAT4 interaction with PAX6. (D) ChIP-WB for validating PAX6 interaction with NFAT4. Error bars indicate standard deviation from three independent experiments, with replicates in each (n=6). *p<0.05, **p<0.01, ***p<0.001, student’s t-test.

## Discussion

The role of lncRNAs in development, differentiation, regulation of lineage specification. and maintenance of pluripotency is well established now with numerous examples reported in the literature. Using high throughput technologies, a large number of lncRNAs have been found to be expressed during neural development and in the brain (Li *et. al*., 2019; Zhao *et. al*., 2020; Roberts *et. al*., 2014; Briggs *et. al*., 2015; Hart & Goff, 2016; Mercer *et. al*., 2008; Derrien *et. al*., 2012; Kadakkuzha *et. al*., 2015; Liau *et. al*., 2021; Lv *et. al*., 2013; Antoniou *et. al*., 2014; Dinger *et. al*., 2008). Owing to their specificity, tight spatial and temporal regulation of expression and functions, lncRNAs have been shown to be more potent indicators of cell types in brain than their protein-coding counterpart, the mRNAs. The roles of many lncRNAs such as Dali, Paupar, Evf2, Rmst, Pnky, to name a few, in regulating neural progenitor proliferation and/or differentiation and gene regulation has already been reported, with new additions every year (Ng *et. al*., 2013; Chalei *et. al*., 2014; Xu *et. al*., 2021; Alammari, 2019; Feng *et. al*., 2006; Stamou *et. al*., 2020; Andersen, 2019; Ramos *et. al*., 2015). A recent study demonstrates that lncRNA SenZfp536 suppresses proliferation of cortical NPCs by cis-regulating the proteincoding gene *Zfp536* via mechanism of chromosomal looping (Tian *et. al*., 2021). While in another study, lncRNA Neat1 was found to induce neuron specific differentiation and to promote migration ability in spinal cord neural stem cells of mice by regulating WNT/β-catenin signaling. In this study, the authors also gave evidence of regulation of lncRNA Neat1 expression by miR-124 (Cui *et. al*., 2019). Study by Lei *et. al*. shows expression level of LncRNA Rik-203 to be increased during neural differentiation. Their data suggest that it facilitates neural differentiation by inhibiting miR-101a-3p’s ability to reduce GSK-3β levels (Zhang *et. al*., 2019 c).

Role of lncRNA Mrhl in development and differentiation has already been established in our previous studies (Arun *et. al*., 2012; Akhade *et. al*., 2014; 2016; Pal *et. al*., 2021). Nuclear restricted Mrhl RNA has a definitive role in differentiation of B type spermatogonial cells into meiotic spermatocytes. Mrhl does so by negatively regulating WNT-signaling pathway, which is activated in differentiated meiotic spermatocytes, in association with p68/DDX5 helicase (Arun *et. al*., 2012). On the other hand, in the context of mESCs, shRNA mediated depletion and subsequent transcriptome analysis have revealed genes related to developmental processes, lineage-specific transcription factors and key networks to be dysregulated along with aberrance in specification of early lineages during differentiation of mESCs (Pal *et. al*., 2021). Interestingly, gene ontology analysis after Mrhl knockdown revealed ‘development process’ category to be enriched as the most important one with maximum number of dysregulated genes, most of them being related to the neuronal lineage. Furthermore, nervous system also emerged as one of the perturbed clusters in gene co-expression analysis. Genome-wide chromatin occupancy studies further predicted important neuronal TFs such as POU3F2 and FOXP2 to be directly regulated at the chromatin level by Mrhl RNA. Our present study validated further that Mrhl is a key molecular player in neuronal lineage development.

Neuronal differentiation is a highly complex process which requires intertwined action of various signaling pathways and transcription factors involved in regulating various aspects of neural stem cells/progenitor proliferation and neurogenesis. The peak of lncRNA Mrhl expression at E14.5 concurrent with the peak of neurogenesis in mouse brain development along with the abrupt inhibition of Mrhl expression from postnatal stage P0 onwards is intriguing. This is also observed in the RA induced neuronal differentiation model system. This observation indicates that Mrhl may have an important role to play in neuronal progenitor specification/maintenance processes during embryonic brain development. Similar observations were also reported by Lv *et.al*. (Lv *et. al*., 2013) where transcriptomics analysis revealed embryonic restricted expression for many lncRNAs. For example, they detected an intronic lncRNA (chr15:66,090,924-66,092,050) which is highly expressed in early embryonic stages and the expression is decreased to basal level from E17.5. As another example, they also identified lncRNA Zfhx2as with increasing expression in E13-E16 peaking at E17 while only basal expression in brain after birth. They concluded that embryonic restricted expression of lncRNAs have regulatory roles in the developing brain (Lv *et. al*., 2013). It would be most interesting to delineate the mechanism of this shutting off of Mrhl expression in the postnatal brain as well as in the maturing neurons in the RA model system.

The mechanisms regulating this developmental stage specific expression of lncRNAs is still not fully understood. However, studies have indicated that chromatin state, including the crosstalk between DNA methylation and histone modification might be one of the mechanisms (Lv *et. al*., 2013; Wu *et. al*., 2010; 2014). It has also been reported that miRNAs, specific DNA sequences as in case of lncRNA known as promoter-associated ncRNAs (pancRNAs), specific location of lncRNA with respect to protein coding genes, and sometimes the transcript itself are involved in modulating the expression of lncRNA in cell and stage specific manner (Cui *et. al*., 2019; Wu *et. al*., 2014; Engreitz *et. al*., 2016). LncRNAs also seem to be regulated by the same set of transcription factors that influence the expression of protein coding genes. A transcriptomic profile study on mouse retinal photo receptor cells, done at six developmental time points in mouse, reveals regulation of ~20% of expressed lncRNAs is under control of the rod differentiation factor, neural retina leucine zipper (NRL), predicting their role in development and differentiation of rod cells (Zelinger *et. al*., 2017). Regulation of lncRNAs by various TFs in cancers has also been reported recently (Sun *et. al*., 2019; Huang *et. al*., 2017). As mentioned before, direct binding and transcriptional regulation of lncRNAs by key mouse ES cell transcription factors, OCT4 and NANOG, have been reported to modulate the pluripotency of ES cells (Mohamed *et. al*., 2010). In our efforts to identify key transcription factors that regulate the differential expression of Mrhl RNA during neuronal development, our bioinformatics analysis revealed the presence of PAX6 DNA binding sites in the upstream region of the Mrhl gene which were also occupied by the PAX6 protein at the chromatin level. PAX6 is an important pioneer transcription factor that has proven role in the regulation of neuronal development (Sansom *et. al*., 2009; Thakurela *et. al*., 2016). PAX6 is also a unique transcription factor in that it does not have a definite cognate DNA binding site but has a set of 10 overlapping DNA binding motifs which function in a very context dependent manner (Xie & Cvekl, 2011). Our extensive analysis presented here has shown that among the 4 PAX6 occupied DNA binding sites, site 1 situated 2904bp upstream of Mrhl TSS, is of relevance for the direct regulation of Mrhl expression in NPCs. Since this PAX6 binding site is far upstream of the TSS, we argued that PAX6 might interact with one of the transcription factor that binds to the proximal promoter region. This assumption was also based on reports wherein it has been shown that PAX6 may not act in isolation but interacts with other transcription factors forming a ternary complex (Kamachi *et. al*., 2001) or with chromatin proteins to regulate promoters and enhancers (Sun *et. al*., 2016). Among the four transcription factors that have binding sites within the proximal promoter region of the Mrhl gene, we found that PAX6 interacts with NFAT4 as monitored by co-immunoprecipitation assays. It is interesting to note that NFAT4 is also neuronal lineage specific transcription factor (Vihma *et. al*., 2016; Wild *et. al*., 2019). Another major observation made in the present study is that WNT signaling is not involved in the regulation of Mrhl gene in RA derived NPCs which is in contrast to its role B-type spermatogonial cells. Thus, it is likely that Mrhl gene is regulated by different sets of transcription factors in a lineage specific manner.

The regulation of lncRNA Mrhl by PAX6/PAX6(5A) through multiple sites of action in NPCs, independently or in probable conjunction with co-factors presents forth a wonderful interplay of lncRNAs and proteins in coordinating cell-specific events. However, recognizing the complexity in lineage specification and interplay between several regulation factors, we also cannot rule out the fact that other TFs or cofactors may also be involved in this mechanism. For example, we found the presence of RARE (retinoic acid response elements) sites on the 3kb upstream promoter region for Mrhl. In the context of the RA induced neuronal model system, this becomes worthy of exploration. Furthermore, it would be interesting to study if this Mrhl-PAX6-NFAT4 interaction dynamics is important for the maintenance of the NPC pool during embryonic brain development as well as probe into other factors that are involved in the aforementioned dynamics. It would be particularly interesting to delineate the chromatin based mechanisms behind regulation of Mrhl by these two TFs in NPCs.

Regulation of spatial and temporal expression of lncRNAs indeed adds to their functional role in context specific manner. Expression of lncRNA Mrhl itself is restricted to testis, liver, kidney & spleen of adult mouse (Nishant *et. al*., 2004; Ganesan & Rao, 2008). It’s expression in mouse ES cells and spermatogonial progenitor cells signifies its role in development (Arun *et. al*., 2012; Akhade *et. al*., 2014; 2016; Pal *et. al*., 2021). The present study has shown that the expression of Mrhl is restricted to embryonic stages of brain and NPCs with minimal basal levels in maturing neurons and postnatal brain. At this juncture, an important question that needs to be addressed in the future is the biological relevance of differential expression of Mrhl RNA during neuronal development. The approach of transcriptome analysis in NPCs depleted of Mrhl should give valuable information about the gene targets of Mrhl RNA in NPCs. We would like to add here that the gene targets of Mrhl lncRNA are quite different in the mouse spermatogonial cells and ES cells (Akhade *et. al*., 2014; Pal *et. al*., 2021) highlighting that the gene targets will be cell type specific and context dependent. Thus, the present study has demonstrated an unexpected and a new role for lncRNA Mrhl in neuronal development. Future studies on the molecular and cellular functions of Mrhl lncRNA in neuronal development should add to the diversity of cellular functions exhibited by this lncRNA.

## Materials and Methods

### Cell lines, plasmids, reagents and oligos

E14TG2a feeder independent mouse ES cell line was a kind gift from Prof. Tapas K. Kundu’s lab (JNCASR, India). mouse ES cells were maintained on 0.2% gelatin (Himedia, TC041) coated dishes and grown in DMEM high glucose (Dulbecco’s modified Eagles’ medium), high glucose (Sigma, D1152), 15% FBS (Gibco, 16000-044), 1X non-essential amino acids (Sigma, M7145), 0.1 mM β-mercaptoethanol and 1X penicillin-streptomycin (Sigma, P4333) supplemented with ESGRO (Merck Millipore). P19EC line was a kind gift from Prof. Kumar Somasundaram’s lab (IISc, India) and grown in in DMEM, 10% FBS, 0.1 mM β-mercaptoethanol and 1X penicillinstreptomycin. HEK293T cell line (ATCC, USA) was maintained in DMEM, 10%FBS and 1X penicillinstreptomycin.

PAX6 shRNA (in the SMART vector inducible lentiviral backbone) clone was obtained from Horizon Discovery, cloneID: V3IMMPGG_11530401.

p3x-FLAG-CMV-10 (Sigma) was used for the generation of PAX6 and PAX6(5A) expression vectors. PAX6 and PAX6(5A) were amplified from cDNA of E14.5 mouse embryonic brains and cloned into the vector.

pGL4.10 (luc2) vector (Promega, E6651) was used for the generation of Mrhl promoter constructs for luciferase assays. 1.25 kb, 1.5 kb or 3.25 kb of the Mrhl promoter (250 bp were taken downstream of the transcription start site in each case) were amplified from genomic DNA and cloned into the vector.

All fine chemicals were obtained from Sigma (unless otherwise mentioned), gelatin was obtained from Himedia and FBS was obtained from Gibco (Performance Plus, US Origin).

Antibodies and oligo sequences have been listed in Supplementary File 2.

### Animal procedures

CD-1 strain of mice (species: *Mus musculus*) were used for isolating embryonic and postnatal brains at indicated stages. For embryonic brain isolation, female mice at age 6-8 weeks were bred with males and were checked for plugs to confirm mating. Day of positive plug was considered as 0.5 dpc. All animal procedures were performed in accordance with the Institutional Animal Ethics Committee of JNCASR, India.

### T ransfection/transduction protocols

Transfections for P19EC and HEK293T cells were performed using Lipofectamine 2000 (Thermo Fisher Scientific, 11668019) as per the manufacturer’s protocol. Transduction of mESCs was performed as per the protocol of Pijnappel *et. al*. (Pijnappel *et. al*., 2013). Briefly, HEK293T cells were transfected with 5 μg PAX6 shRNA plasmid, 2.5 μg pSPAX2, 1.75 μg pVSVG and 0.75 μg pRev to generate the viral particles. The media containing viral particles was harvested 48 hours after transfection. A second round of viral particles was collected after an additional 24 hours. The viral supernatant was mixed with 8 μg/ml DEAE-dextran and 1000 units/ml ESGRO and added directly to the E14TG2a cells. Transduction was performed for 24 hours with the first round of viral particles and an additional 24 hours with the second round of viral particles. The transduced cells were then subjected to puromycin selection (1.5 μg/ml puromycin) for a week.

### Neuronal differentiation of mESCs

The protocol described by Bibel *et. al*. (Bibel *et. al*., 2007) was followed for retinoic acid mediated neuronal differentiation of mESCs. 4X10^6^ cells were plated onto 100 mm bacteriological grade dishes (Greiner Bio-One, 633102) in differentiation medium containing DMEM, 10% FBS, 0.1 mM β-mercaptoethanol and 1X penicillin-streptomycin and allowed to grow for 4 days. Retinoic acid [RA (Sigma, R2625)] or dimethyl sulfoxide [(DMSO), Sigma, D2650] was added to the EBs on the 4^th^ day at a final concentration of 5 μM and allowed to grow for an additional 4 days. After a total of 8 days of differentiation (4-/4+ RA), the RA or vehicle DMSO treated EBs were harvested by gravity precipitation for cryosectioning. Alternatively, RA treated EBs were trypsinized to obtain NPCs with 0.05% freshly made trypsin for 3 minutes at 37°C, dissociated and filtered through 40 μm filters (Falcon, 352340). NPCs were collected by centrifugation at 1150 rpm for 5 minutes.

For the generation of early and maturing neurons, plates or coverslips were coated with 100 μg/ml poly-D-lysine (Merck Millipore, A-003-E) at 37°C overnight. Next day, plates were washed thoroughly three times with deionized sterile water and dried. The plates were then coated with laminin solution (Roche, 11243217001) for 2 hours at 37°C and kept ready for use. The cell pellet obtained above was resuspended in N2 medium containing DMEM/F12 (Sigma, D2906), 1X N2 (Thermo Fisher Scientific, 17502048) and 1X penicillin-streptomycin and plated at a density of 90,000 cells/cm^2^. N2 medium was changed 2 hours after plating and then again after 24 hours. For further differentiation, medium was changed after an additional 24 hours to N2B27 medium containing DMEM/F12, 1X N2, 1X B27 (Thermo Fisher Scientific, 17504044) and 1X penicillin-streptomycin. Cells were harvested at different time points for appropriate analyses.

### PAX6 knockdown and Mrhl expression

PAX6 shRNA clone was used to generate stable inducible mESC line. Protocol for retinoic acid mediated neuronal differentiation of mESCs as described above was followed. PAX6 knockdown was induced by addition of different concentrations of doxycycline (0, 2, 3, 3.5 & 4 μg/ml) from day 3 onwards, every day. RA was added on 4^th^ day and EBs were harvested on day 5^th^. RNA was isolated from collected cells and qPCR was performed as described later, for expression level of PAX6 and Mrhl.

### NOTCH Inhibition studies

EBs were treated with NOTCH inhibitor DAPT (N-[N-(3,5-Difluorophenacetyl)-L-alanyl]-S-phenylglycine t-butyl ester, Sigma D5942) at a concentration of 10 μM on the 4^th^ day of differentiation along with RA. EBs were harvested on the 5^th^ day and processed directly for qPCR as described later.

### Preparation of NPC smears

EBs were trypsinized and filtered as described earlier and cross-linked with 1% formaldehyde for 8 minutes at room temperature. Formaldehyde was quenched using 1/7^th^ volume of 1M glycine for 5 minutes at room temperature and cells were collected by centrifugation at 1500 rpm for 5 minutes at 4°C. The cells were washed 3 times with ice cold 1X PBS and collected by centrifugation each time. A 10 μl aliquot of the cell suspension in 1X PBS was taken on a slide, covered with a coverslip and dipped in liquid N_2_ to fix the cells on the slide (aliquot volume can be determined empirically to obtain desired cell density per aliquot). The cover slip was removed immediately thereafter with the help of scalpel blade. The slides were hydrated in 1X PBS for 3-5 minutes and dehydrated in graded ethanol from 50%, 75%, 95% to 100%. The slides can be stored in 100% ethanol at this point. Before proceeding for RNA FISH or IF, slides were rehydrated in the reverse order of ethanol series with a final rehydration step in 1X PBS.

### RNA fluorescent in situ hybridization (FISH) and immunofluorescence (IF)

RNA FISH followed by IF on EBs, NPCs or brain tissues was performed as per the protocol of de Planell-Saguer *et. al*. (de Planell-Saguer *et. al*., 2010). The probes used for RNA FISH were Cy5 labelled locked nucleic acid probes procured from Exiqon (reported in Arun *et. al*.) (Arun *et. al*., 2012).

#### Fixation and permeabilization

Cells were either grown on coverslips or cell smears were prepared as described above. EBs or brain tissues were fixed in 4% paraformaldehyde for 7-8 hours at 4°C followed by equilibration in 20% sucrose solution overnight and embedded in tissue freezing medium (Leica, 14020108926). The embedded tissues were then cryo-sectioned and collected on Superfrost slides (Fisher Scientific, 12-550-15). For cells, a brief wash was given with 1X PBS (phosphate buffered saline, pH 7.4) followed by fixation with 2% formaldehyde for 10 minutes at room temperature. The cells were then washed with 1X PBS three times for 1 minute each and permeabilization buffer (1X PBS, 0.5% Triton X-100) was added for 5 minutes and incubated at 4°C. The permeabilization buffer was removed and cells were washed briefly with 1X PBS for three times at room temperature. For tissue sections, antigen retrieval was performed by boiling the sections in 0.01M citrate buffer (pH 6) for 10 minutes. The sections were allowed to cool, washed in distilled water three times for 5 minutes each and then in 1X PBS for 5 minutes, each time with gentle shaking.

#### Hybridization

The samples were then blocked in prehybridization buffer [3% BSA, 4X SSC (saline sodium citrate, pH 7)] for 40 minutes at 50°C. Hybridization (Mrhl probes tagged with Cy5, final concentration 95nM) was performed with prewarmed hybridization buffer (10% dextran sulphate in 4X SSC) for 1 hour at 50°C. After hybridization, slides were washed four times for 6 minutes each with wash buffer I (4X SSC, 0.1% Tween-20) at 50°C followed by two washes with wash buffer II (2X SSC) for 6 minutes each at 50°C. The samples were then washed with wash buffer III (1X SSC) once for 5 minutes at 50°C followed by one wash with 1X PBS at room temperature. For tissue sections, all washes were performed as mentioned above with a time of 4 minutes for buffers I-III. The samples were then processed for IF.

#### IF

Samples were blocked with IF Blocking buffer (4% BSA, 1X PBS) for 1 hour at 4 °C. Primary antibody solution (2% B SA, 1X PBS) was prepared containing the appropriate dilution of desired primary antibody and the samples were incubated in it for 12 hours at 4°C. Next day, the samples were washed with IF wash buffer (0.2% BSA, 1X PBS) three times for 5 minutes each with gentle shaking. The samples were incubated in secondary antibody for 45 minutes at room temperature and washed with 1X PBS three times for 10 minutes each with gentle shaking. The samples were finally mounted in mounting medium containing glycerol and DAPI.

### Chromatin IP (ChIP)

ChIP was performed according to Cotney and Noonan’s protocol (Cotney & Noonan, 2015). Protein A or protein G Dynabeads (Thermo Fisher Scientific, 10001D, 10003D) was used for rabbit or mouse antibodies respectively. For co-ChIP, same protocol was followed and whole sample after IP was processed for western blotting.

#### Preparation of antibody beads

25 μl dynabeads were used for each ChIP reaction. The beads were first washed with 1 ml of bead binding buffer (1X PBS, 0.2% Tween-20) and resuspended in 200 μl of the buffer per reaction. H3K4me3 (2 μg) or Pax6 (4 μg) antibodies or their isotype controls were added to the beads in separate tubes and incubated for ~16 hours at 4°C on an end-to-end rotor at 10 rpm. The next day, beads were washed with 1ml of bead binding buffer followed by 1ml of dilution buffer (0.01% SDS, 1.1% Triton X-100, 1.2 mM EDTA, 16.7 mM Tris-HCl pH 8.1, 167 mM NaCl). The beads were then resuspended in 25 μl dilution buffer per ChIP reaction and stored at 4°C for further use.

#### Chromatin extraction and quantification

Brain tissues from E14.5 dpc embryos or EBs were harvested in serum-free DMEM. Brain samples were minced with a scalpel blade and pipetted a few times whereas EBs were trypsinized and filtered as described earlier. The samples were then subjected to cross-linking using 1% formaldehyde (Sigma, F8775) for 10 minutes at room temperature following which samples were quenched with glycine at a final concentration of 0.125M for 10 minutes at room temperature. The samples were then centrifuged at 2,000 g for 10 minutes at 4°C and washed twice with 1X ice-cold PBS. Finally, the pellets were either flash frozen in liquid N_2_ and stored in −80°C or processed for chromatin extraction.

The cross-linked pellet was resuspended in six volumes of ice-cold cell lysis buffer (50 mM Tris-HCl pH 8.0, 140 mM NaCl, 1mM EDTA, 10% glycerol, 0.5% NP-40, 0.25% Triton X100) supplemented with 1X mammalian protease inhibitor cocktail and incubated on ice for 20 minutes. For brain samples, the pellets were homogenized once after addition of lysis buffer with 30-40 strokes (BioSpec tissue tearor, 985370) and once after the incubation period was over. The nuclei were then harvested by centrifugation at 2,000 g for 5 minutes at 4°C. The supernatant was removed, the nuclei were resuspended in five volumes of ice-cold nuclear lysis buffer (10 mM Tris-HCl pH 8.0, 1 mM EDTA, 0.5 mM EGTA, 0.5% SDS) supplemented with 1X mammalian protease inhibitor cocktail and incubated on ice for 20 minutes. The samples were then sonicated in Bioruptor (Diagenode, UCD-200) for 35 cycles (at pulses of 30sec on and 30sec off) and centrifuged at 16,000 g for 10 minutes at 4°C to remove insoluble material. The centrifuged samples were then transferred to fresh tubes, aliquoted as per requirement, flash frozen in liquid N_2_ and stored in −80°C.

For monitoring sonication efficiency and DNA quantification, 10 μl aliquots was kept aside, diluted in 10 μl of TE buffer and treated with 10 μg of RNaseA (Sigma, R6513) for 30 minutes at 37°C followed by 20 μg of proteinase K (Thermo Fisher Scientific, EO0491) for 1 hour at 55 °C. The aliquots were then subjected to reverse cross-linking for 5 minutes at 95 °C, allowed to cool to room temperature slowly and analyzed on 1% agarose gel for sonication efficiency or subjected to DNA isolation by phenol-chloroform method for quantification.

#### Immunoprecipitation

Approximately 10-25 μg of chromatin was diluted with dilution buffer supplemented with 1X mammalian protease inhibitor cocktail to reduce the SDS concentration to <0.1% and achieve a final volume of 450 μl. 5% of the dilution was stored as input at 4°C. 25 μl of antibody or isotype control beads were added to the chromatin dilution and incubated for 12-16 hours at 4°C. Next day, the beads were washed with 1 ml wash buffer (100 mM Tris-HCl pH 8.0, 500 mM LiCl, 1% NP-40, 1% deoxycholic acid) supplemented with 1X mammalian protease inhibitor cocktail for 5 times at room temperature. The beads were given a final wash with 1 ml of TE buffer and resuspended in 85 μl of elution buffer (50 mM Tris-HCl pH 8.0, 10 mM EDTA, 1% SDS). Elution was performed twice for 10 minutes each at 65 °C under constant agitation in a thermo mixer. All ChIP and input samples (volume made upto 170 μl with dilution buffer for input samples) were subjected to reverse crosslinking for 12 hours at 65°C.

#### Chromatin purification and analysis

Next day, all samples were treated with 10 μg of RNAse A for 1 hour at 37°C followed by 200 μg proteinase K for 2 hours at 55 °C and DNA was extracted by the phenol-chloroform method. The DNA was precipitated with 1/10^th^ volume 3M sodium acetate (ph 5.2), 3 volumes of 100% ethanol and glycogen at a final concentration of 0.5 μg/μl overnight at −20°C. Next day, DNA was pelleted, washed with 75% ethanol, dried, dissolved in sterile deionized water and subjected to qRT-PCR analysis.

### Purification of PAX6 and PAX6(5A) proteins

p3X-FLAG-CMV-10-Pax6 and -Pax6(5a) constructs were transfected into HEK293T cells. 24 h after transfection, cells were lysed in five volumes of FLAG lysis buffer (50 mM Tris-HCl pH 7.4, 150 mM NaCl, 1 mM EDTA, 1% Triton X-100) supplemented with 1X mammalian protease inhibitor cocktail. To 500 μl of supernatant containing proteins, 30 μl M2 agarose beads (Sigma, A220) were added and incubated for 3 h at 4°C. The beads were then washed in 1 ml of FLAG wash buffer (50 mM Tris-HCl pH 7.4, 150 mM NaCl). The FLAG tagged proteins were then eluted in the FLAG wash buffer containing a final concentration of 500 ng/μl of FLAG peptide (Sigma, F4799).

### Electrophoretic Mobility Shift Assay (EMSA)

#### End labelling of probe and purification

Oligos were designed such that they harboured the PAX6/PAX6(5A) motifs on Mrhl promoter. All forward oligos were subjected to end labelling with γ-32P-ATP in polynucleotide kinase reaction buffer [PNK buffer in a final concentration of 1X, 40pmol oligo, 20 units PNK enzyme (NEB, M0201S), 30 μCi γ-32P-ATP, volume made upto 50 μl with nuclease free water] for 30 minutes at 37°C. The enzyme was heat inactivated for 20 minutes at 65 °C. The oligos were purified by the phenol-chloroform method and precipitated in 1/10^th^ volume of 3M sodium acetate, ph 5.2, 3 volumes of 100% ethanol and 10 μg yeast tRNA for 1 hour at −80°C. The oligos were pelleted by centrifugation at 12,800 rpm for 10 minutes, washed in 75% ethanol and dried. They were then dissolved in 1X annealing buffer (10 mM Tris-HCl ph 8.0, 20 mM NaCl). To each of the forward oligos, three times excess of the reverse oligos were added, heated for 10 minutes at 95 °C and allowed to cool slowly over several hours to overnight at room temperature. The annealed probes were purified using sephadex C-50 columns. Glass wool was packed near the tip of a 1 ml syringe till 0.1 ml and the syringe was packed with C-50 beads slurry till 1 ml with centrifugation at 1800 rpm for 3 minutes at 4°C. A final centrifugation was performed for the packed beads to remove all water before loading the annealed probes onto the column. The annealed probes were collected again by centrifugation and their activity was noted as counts per minute (cpm) in a scintillation counter. Unlabelled forward oligos were also annealed to their corresponding reverse oligos and purified in a similar manner for competition assay.

#### Binding reaction and electrophoresis

The purified proteins were allowed to bind to the oligos in a reaction mix containing 5X EMSA Buffer (60 mM Hepes-KOH ph 7.9, 300 mM KCl, 15 mM MgCl_2_, 2.5 mM DTT, 20% w/v Ficoll 400, 1 mg/ml BSA, 1X mammalian protease inhibitor cocktail added just before use) to a final concentration of 1X, PAX6 or PAX6 (5A) purified proteins (100ng to 400ng in increasing concentrations), 400 ng/μl salmon sperm DNA (Sigma, D9156) and y-32P-ATP labelled double stranded oligo (20,000 cpm). The volume was made upto 12.5 μl with nuclease free water. The reaction was incubated for 40 minutes at 37°C. For competition assay, 125X molar excess of unlabelled double stranded probe was used. After incubation, loading dye was added to the samples (containing bromophenol blue, xylene cyanol and without SDS) and the samples were run on a 5% native gel (40% acrylamide stock in 30:08 ratio of acrylamide to bis-acrylamide) prepared in 0.5X TBE (10X TBE stock: 1.8M tris base, 90mM boric acid, 2.5mM EDTA) at 150V at 4°C. The gel was then dried and exposed to X-ray film for 24 hours.

### Luciferase and β-galactosidase Assay

P19EC cells were transfected with 2 μg of pGL4-Mrhl promoter constructs, 200 ng of PAX6 or PAX6(5A) expression plasmids and 50 ng of β-gal plasmid as an internal control. Cells were harvested 24 h post transfection and luciferase assay was performed as per the manufacturer’s protocol (Promega, E2610). Readings for luminescence were taken in luminometer (Berthold, Sirius L).

The β-gal assay as performed as per Uchil *et. al*. (Uchil *et. al*., 2017). Briefly, 30 μl of the cell extract was mixed with 3 μl 100X Mg^+2^ solution (0.1 M MgCl_2_, 4.5 M beta-mercaptoethanol), 66 μl 1X ONPG (4 mg/ml ONPG in 0.1 M phosphate buffer pH 7.5) and 201 μl 0.1 M sodium phosphate, pH 7.5. The reactions were incubated for 30 minutes to a few hours at 37°C until a faint yellow color developed. The reactions were stopped by adding 500 μl 1 M Na_2_CO_3_ and the absorbance was recorded at 420 nm in a spectrophotometer.

### RNA isolation, qPCR and Western Blotting

Total RNA was isolated from cells or tissues using RNAiso Plus (Takara Bio) for analysis by qRT-PCR as per the manufacturer’s protocol. Real-time PCR was performed using SyBr green mix (Takara) in Realtime PCR machine (BioRad CFX96).

Cells or tissues were lysed in five times volume of RIPA buffer (150 mM sodium chloride, 1.0% NP-40 or Triton X-100, 0.5% sodium deoxycholate, 0.1% SDS, 50 mM Tris-HCl pH 8.0, 1 mM EDTA, 0.5 mM EGTA) supplemented with 1 mM PMSF and 1X mammalian protease inhibitor cocktail for 15 minutes on ice with occasional vortexing. Protein concentration was estimated using Bradford reagent. Samples were resolved on 10-12% SDS-PAGE gel, transferred onto nitrocellulose membrane, incubated with primary and secondary antibodies and analyzed using chemiluminescence (Millipore, Luminata Forte, WBLUF0100) and exposure to X-ray films.

### FIMO analysis

3 kb promoter sequence of Mrhl (upstream from its TSS) was taken along with the motifs for PAX6 binding as described in a report by Xie and Cvekl (Xie and Cvekl, 2011). FIMO (Find Individual Motif Occurrences) (Grant *et. al*., 2011) scanning was then performed with a default cutoff value of 1E-04.

### Promoter analysis

GPMiner (Lee *et. al*., 2012) and JASPAR (Mathelier *et. al*., 2014) programs were used to analyze 3kb upstream of transcription start site (TSS) of Mrhl. SnapGene software was used to map the binding sites of various TFs on the promoter region.

## Acknowledgements

We thank Suma B.S of the Confocal Imaging Facility, Dr. R.G Prakash of the Animal Facility and Anitha G. of the Sequencing Facility at JNCASR, India. M.R.S Rao acknowledges Department of Science and technology, Govt of India for SERB Distinguished Fellowship and SERB-YOS (Year of Science Chair Professorship). Debosree Pal thanks University Grants Commission, Govt. of India and JNCASR, India for her PhD fellowship. Sangeeta Dutta thanks Department of Biotechnology, Govt. of India for her postdoctoral fellowship.

## Author CRediT Statement

**Conceptualization:** M.R.S., D.P. **Methodology:** D.P., S.D., D.P.I., M.R.S. **Software:** U.B. **Validation:** S.D. **Formal Analysis:** D.P., S.D., D.P.I., U.B. **Investigation:** D.P., S.D., D.P.I., U.B. **Resources:** M.R.S. **Data Curation:** D.P., U.B. **Writing - Original draft and preparation:** D.P., S.D. **Writing - review and editing:** M.R.S. **Visualization:** D.P., M.R.S. **Supervision:** M.R.S. **Project administration:** M.R.S. **Funding acquisition:** M.R.S.

---

**Figure S1.**
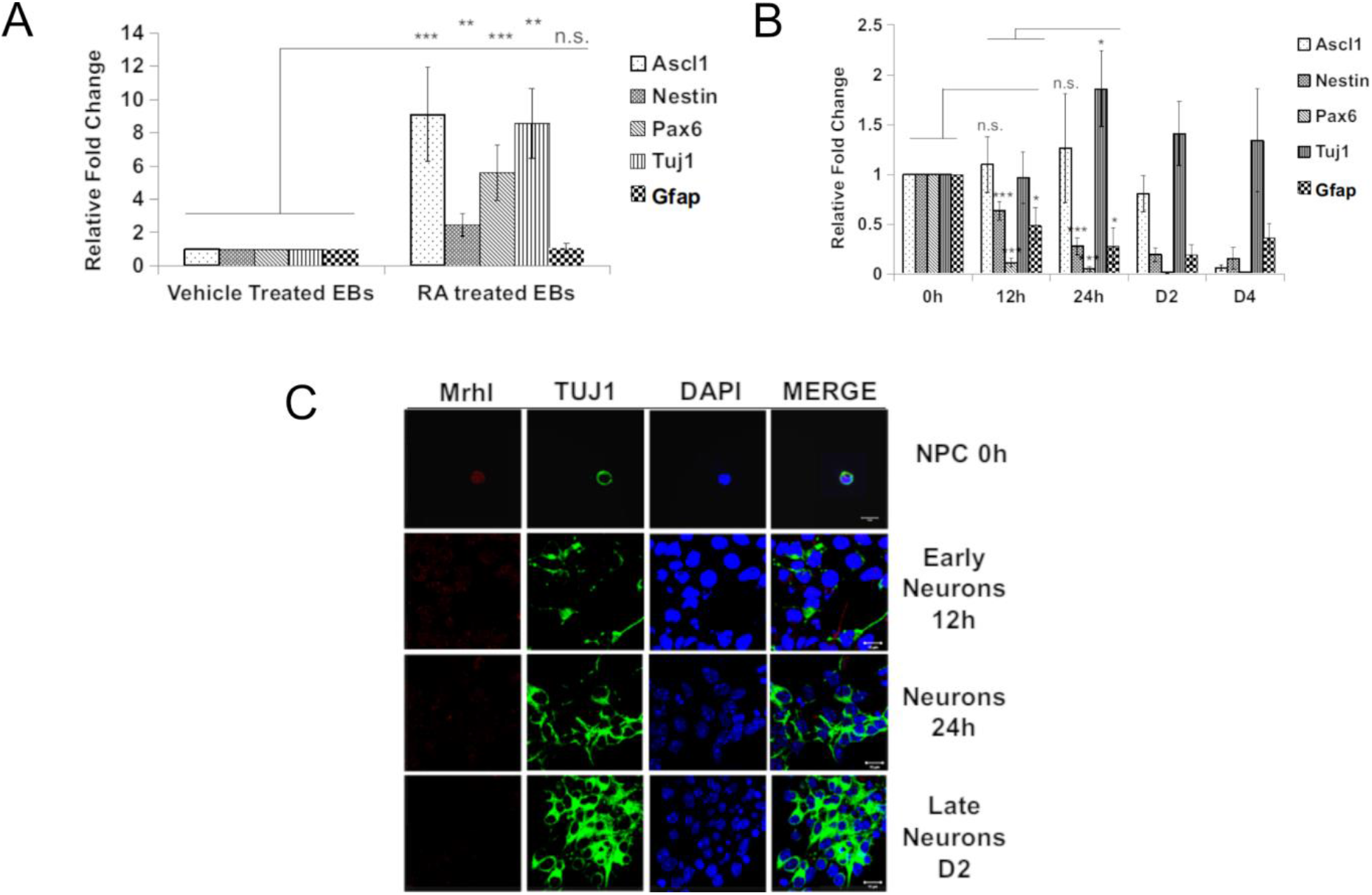
(C) qPCR analysis of various markers for neuronal lineage in RA treated versus vehicle treated EBs. Ascl1 is a neuronal lineage marker. Nestin and Pax6 are NPC markers. Tuj1 is a maturing neuron marker. Gfap is an astrocyte lineage marker. (B) qPCR analysis of various markers for NPCs and neurons at different hours of differentiation show reduction in NPC markers and increase in neuron marker. (E) RNA FISH for Mrhl followed by IF for TUJ1 in neurons. Error bars indicate standard deviation from three independent experiments, with replicates in each (n=6). *p<0.05, **p<0.01, ***p<0.001, student’s t-test; Scale bar = 10 μm.

**Figure S2.**
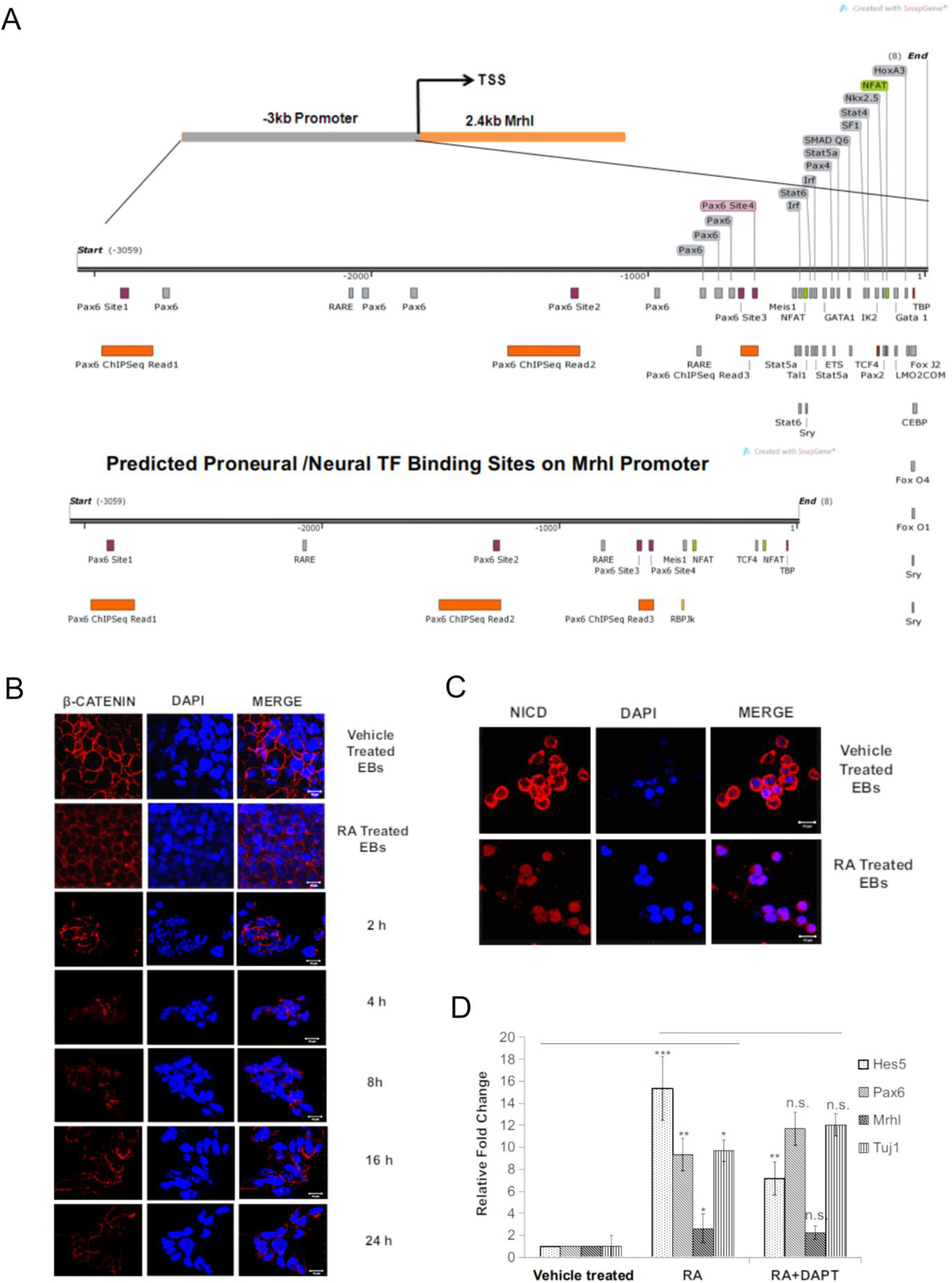
(A) GPMiner and JASPAR analysis for predicted TF binding sites on Mrhl 3kb promoter region; created by SnapGene. (B) IF for β-CATENIN in RA treated versus vehicle treated EBs and different time points of neuronal differentiation. (C) IF for NICD in RA versus vehicle treated EBs. (D) qRT-PCR analysis for genes upon DAPT treatment. Error bars indicate standard deviation from three independent experiments, with replicates in each (n=6). *p<0.05, **p<0.01, ***p<0.001, student’s t-test; Scale bar = 10 μm.

**Figure S3.**
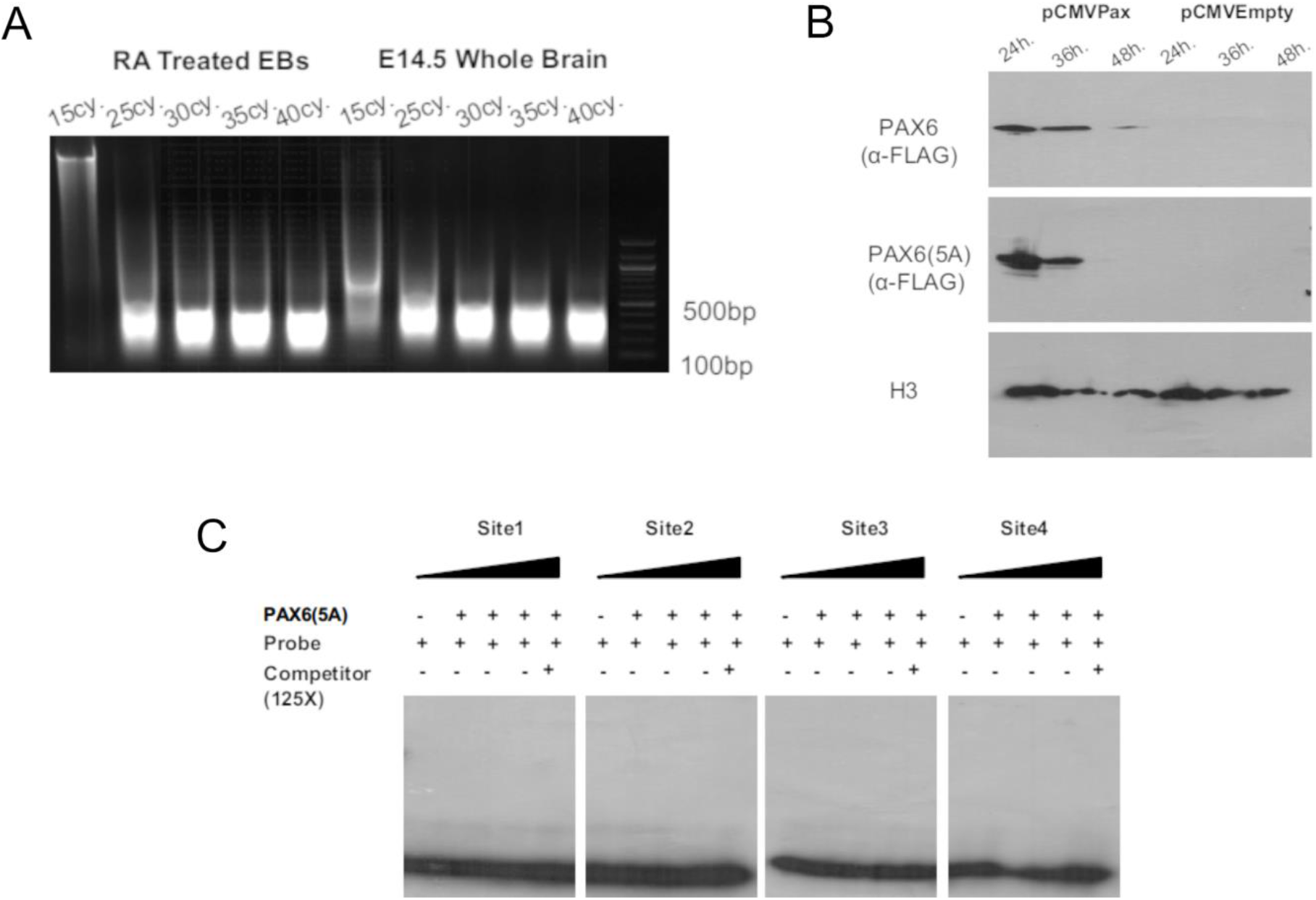
(A) Sonication pattern for chromatin isolated from RA treated EBs and E14.5 whole brain. (B) Expression and validation of PAX6 and PAX6(5A) by western blot. (C) EMSA for PAX6(5A) binding to Mrhl promoter sites 1-4.

**Figure S4.**
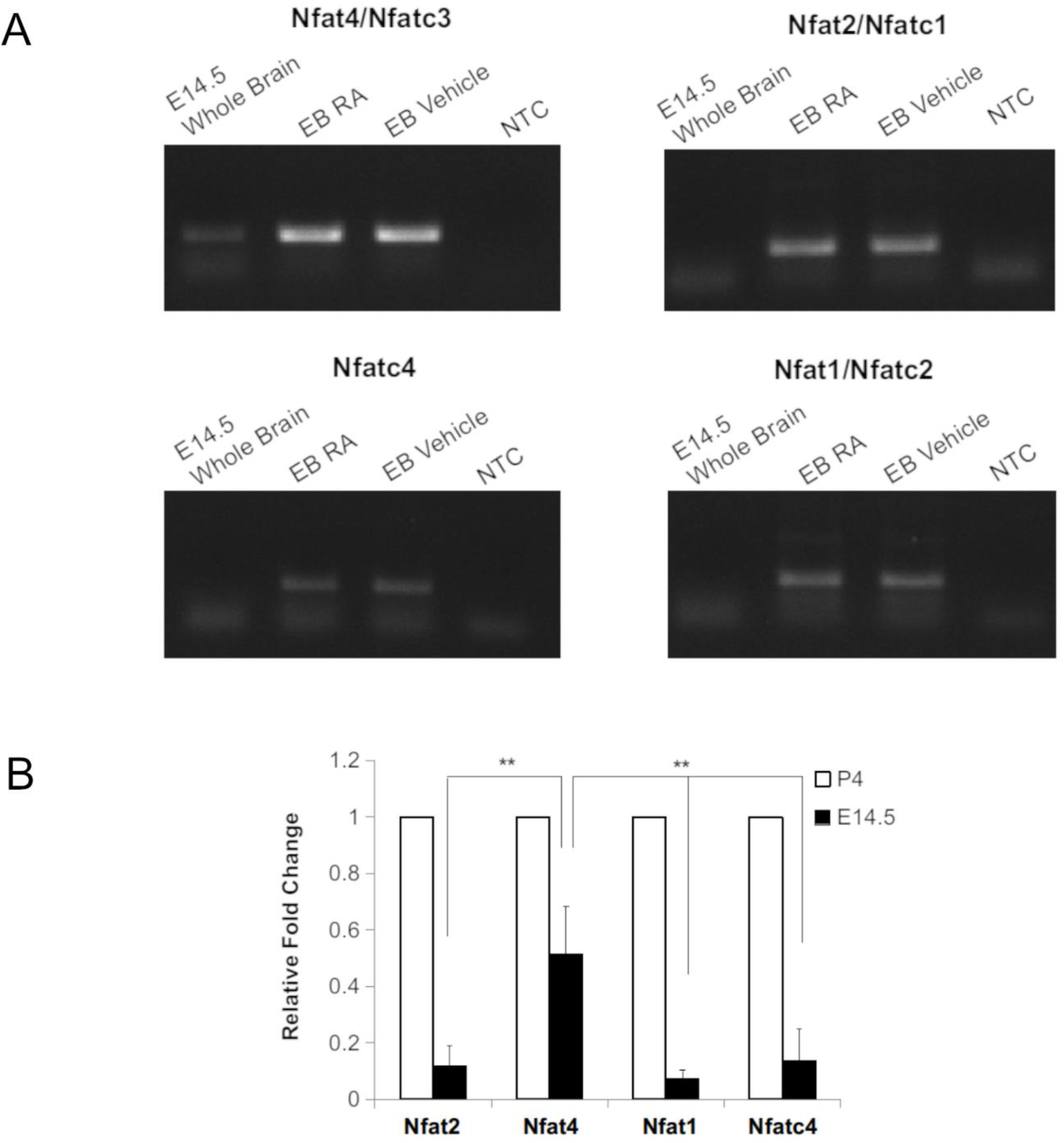
(A) Expression levels of isoforms of NFAT in E14.5 whole brain and EBs; NTC:no template control. (B) qPCR showing NFAT4 expression amongst all isoforms in E14.5 brain as compared to P4 brains. Error bars indicate standard deviation from three independent experiments, with replicates in each (n=6). *p<0.05, **p<0.01, ***p<0.001, student’s t-test.

## Supplementary File 1 (a SnapGene file)

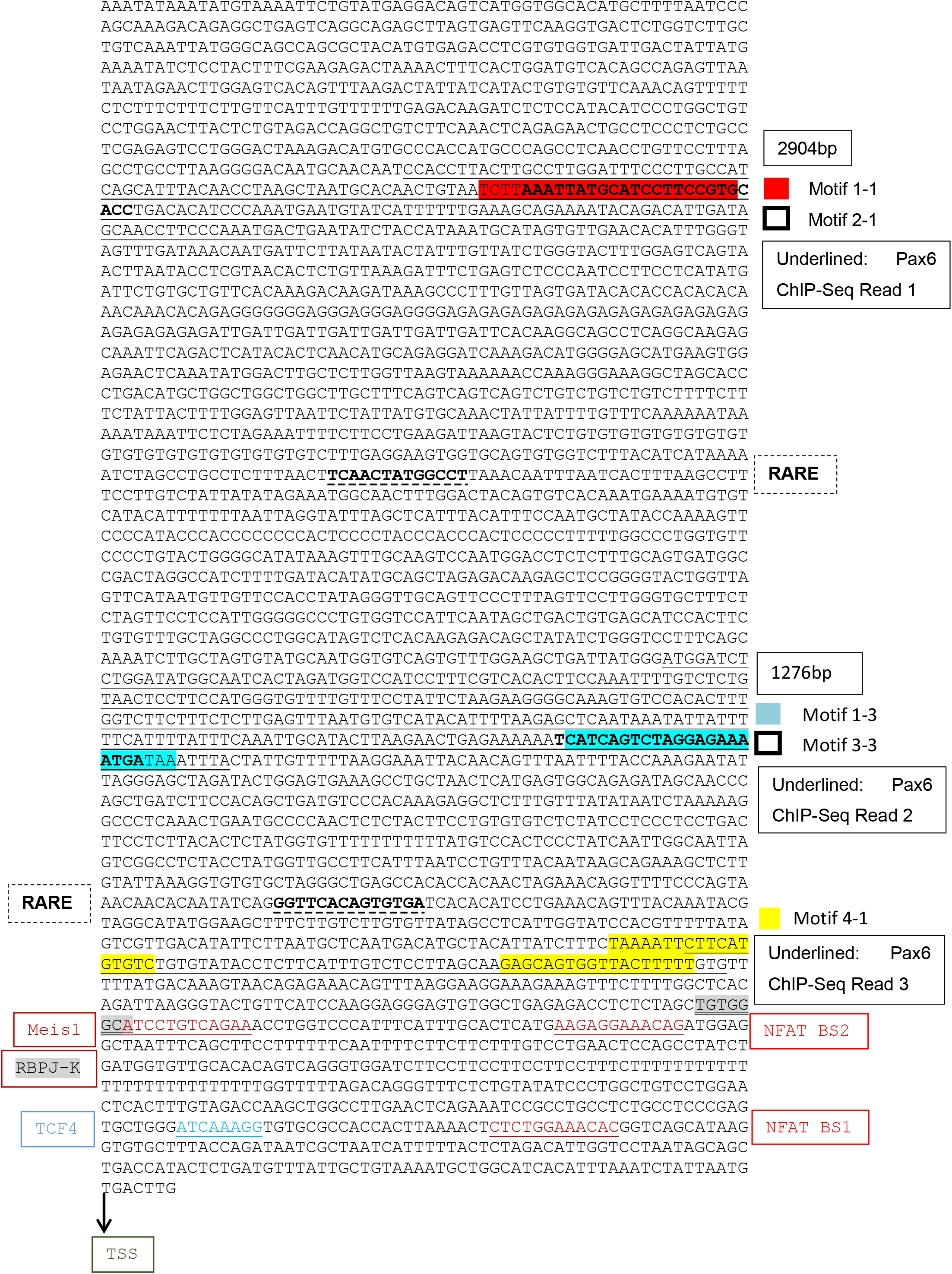

## Supplementary File 2

### Antibodies

| Name of antibody | Company | Catalog No. |
| --- | --- | --- |
| Anti-H3 | Abcam | ab46765 |
| Anti-NESTIN | Abcam | ab11306 |
| Anti-TUJ1 | Abcam | ab14545 |
| Anti-H3K4me3 | Abcam | ab12209 |
| Anti-NICD | Abcam | ab8915 |
| Anti-PAX6 | Abcam | ab5790 |
| Anti-β-catenin | Abcam | Ab6302 |
| Anti-MEIS1 | Millipore | ABE2864 |
| Anti-TCF4 | Cell signaling technology | 2569 |
| Anti-NFAT4/NFATc3 | Invitrogen | PA5-79734 |

### Primer and Oligo Sequences

#### qRT-PCR primers

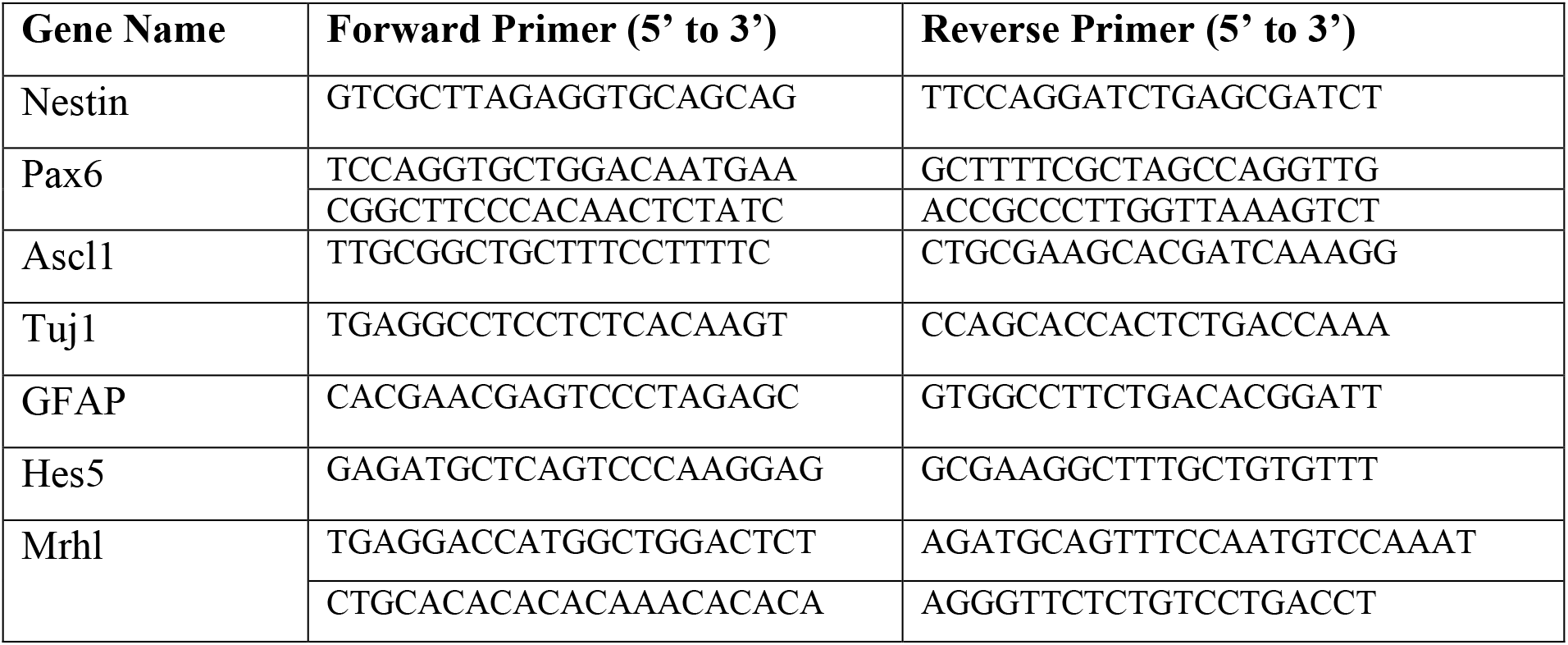
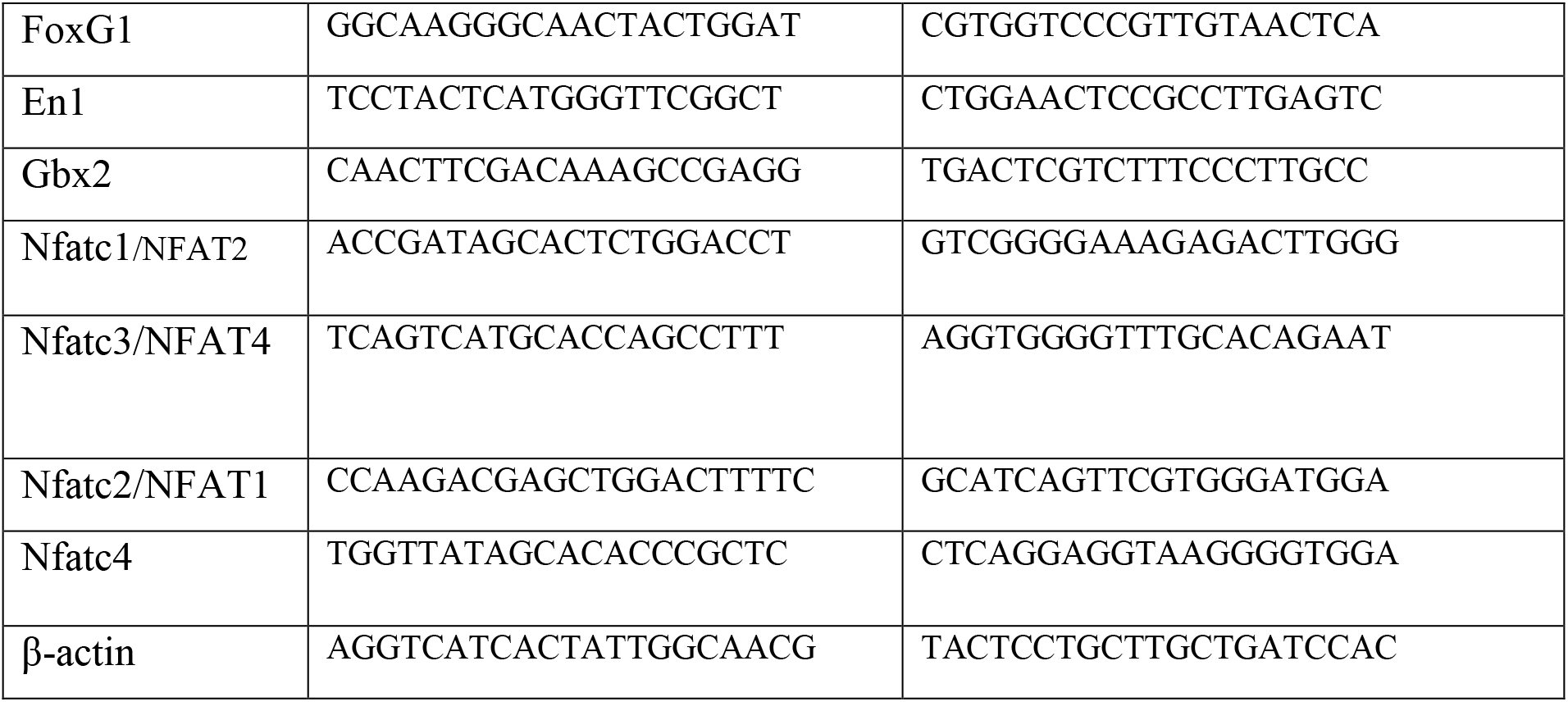

### ChIP primers

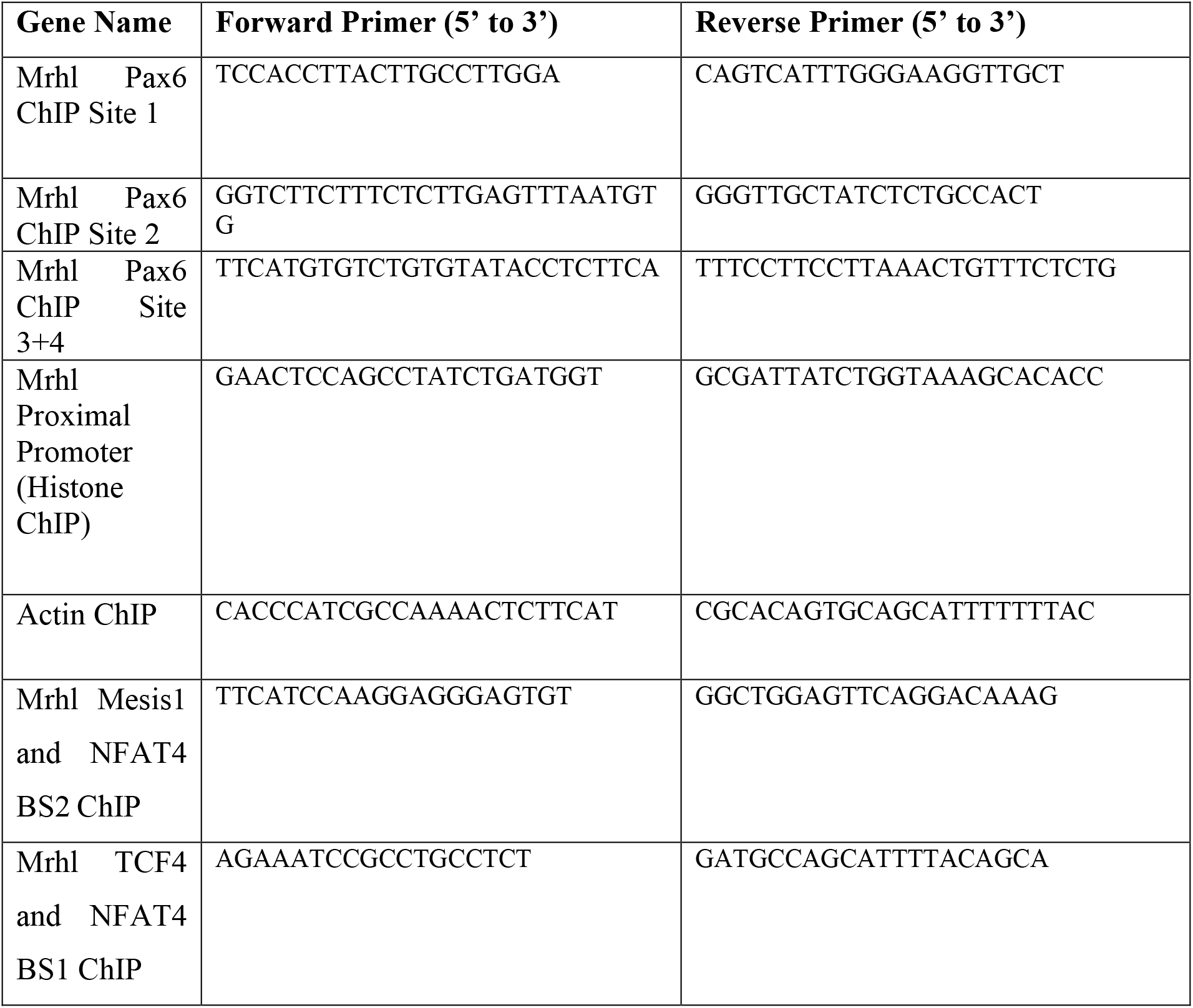

### Cloning primers

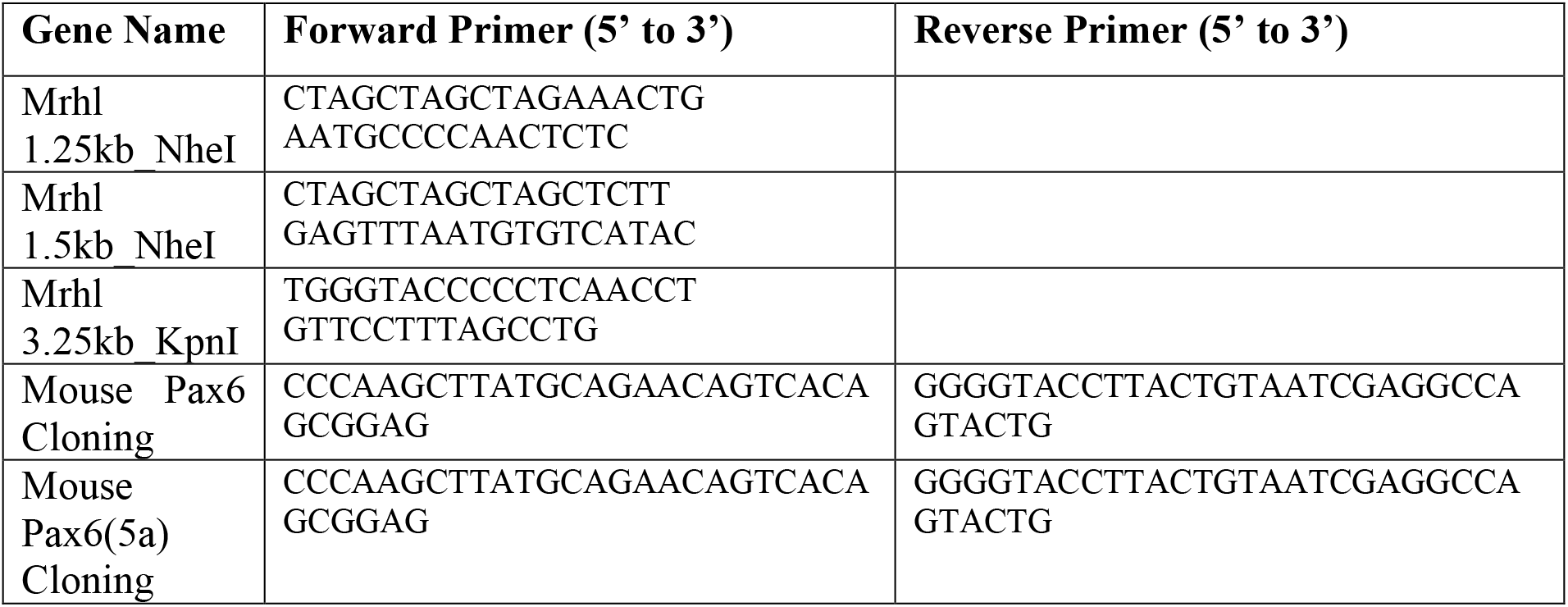

### EMSA oligos (5’ to 3’)

**Site 1:**

Probe 1_F: CTGTAATCTTAAATTATGCATCCTTCCGTGCACCTGACAC
Probe 1_R: GTGTCAGGTGCACGGAAGGATGCATAATTTAAGATTACAG

**Site 2:**

Probe 2_F: AGAAAAAATCATCAGTCTAGGAGAAAATGATAAATTTACT
Probe2_R: AGTAAATTTATCATTTTCTCCTAGACTGATGATTTTTTCT

**Site 3:**

Probe 3_F: CATTATCTTTCTAAAATTCTTCATGTGTCTGTGTATACCTC
Probe 3_R: GAGGTATACACAGACACATGAAGAATTTTAGAAAGATAATG

**Site 4:**

Probe 4_F: TCTCCTTAGCAAGAGCAGTGGTTACTTTTTGTGTTTTTAT
Probe 4_R: ATAAAAACACAAAAAGTAACCACTGCTCTTGCTAAGGAGA

**Positive control for Pax6:**

P6CON_F: GGATGCAATTTCACGCATGAGTGCCTCGAGGGATCCACGTCGA
P6CON_R: TCGACGTGGATCCCTCGAGGCACTCATGCGTGAAATTGCATCC

## References

1. Akhade, V.S., Arun, G., Donakonda, S. and Satyanarayana Rao, M.R. (2014). Genome wide chromatin occupancy of mrhl RNA and its role in gene regulation in mouse spermatogonial cells. RNA biology, 11(10), pp.1262–1279.

2. Akhade, V.S., Dighe, S.N., Kataruka, S. and Rao, M.R.S. (2016). Mechanism of Wnt signaling induced down regulation of mrhl long non-coding RNA in mouse spermatogonial cells. Nucleic acids research, 44(1), pp.387–401.

3. Akhade, V.S., Pal, D. and Kanduri, C. (2017). Long noncoding RNA: genome organization and mechanism of action. Long Non Coding RNA Biology, pp.47–74.

4. Alammari, F. (2019). KAP1-Paupar lncRNA chromatin regulatory complex controls subventricular zone neurogenesis. PhD thesis, University of Oxford, Oxford, United Kingdom.

5. Andersen, R.E. (2019). The Novel Long Noncoding RNA Pnky Regulates Neurogenesis and Neural Stem Cell Maintenance In Vivo. PhD thesis, University of California, San Francisco, USA.

6. Antoniou, D., Stergiopoulos, A. and Politis, P.K. (2014). Recent advances in the involvement of long non-coding RNAs in neural stem cell biology and brain pathophysiology. Frontiers in physiology, 5, p.155.

7. Arun, G., Akhade, V.S., Donakonda, S. and Rao, M.R.S. (2012). mrhl RNA, a long noncoding RNA, negatively regulates Wnt signaling through its protein partner Ddx5/p68 in mouse spermatogonial cells. Molecular and cellular biology, 32(15), p.3140.

8. Bibel, M., Richter, J., Lacroix, E., and Barde, Y. A. (2007). Generation of a defined and uniform population of CNS progenitors and neurons from mouse embryonic stem cells. Nature protocols, 2(5), 1034.

9. Bouchard, M., Grote, D., Craven, S. E., Sun, Q., Steinlein, P., and Busslinger, M. (2005). Identification of Pax2-regulated genes by expression profiling of the midhindbrain organizer region. Development, 132(11), 2633–2643.

10. Bouilloux, F., Thireau, J., Venteo, S., Farah, C., Karam, S., Dauvilliers, Y., Valmier, J., Copeland, NG., Jenkins, NA., Richard, S., et. al.(2016). Loss of the transcription factor Meis1 prevents sympathetic neurons target-field innervation and increases susceptibility to sudden cardiac death. Elife, 5, e11627.

11. Briggs, J.A., Wolvetang, E.J., Mattick, J.S., Rinn, J.L. and Barry, G. (2015). Mechanisms of long non-coding RNAs in mammalian nervous system development, plasticity, disease, and evolution. Neuron, 88(5), pp.861–877.

12. Chalei, V., Sansom, S.N., Kong, L., Lee, S., Montiel, J.F., Vance, K.W. and Ponting, C.P. (2014). The long non-coding RNA Dali is an epigenetic regulator of neural differentiation. elife, 3, p.e04530.

13. Chen, Y. and Tergaonkar, V. (2020). LncRNAs: Master Regulators in Disease and Cancer. Proceedings of the Singapore National Academy of Science, 14(02), pp.79–89.

14. Chenn, A., and Walsh, C. A. (2002). Regulation of cerebral cortical size by control of cell cycle exit in neural precursors. Science, 297(5580), 365–369.

15. Choudhury, S.R., Dutta, S., Bhaduri, U. and Rao, S.M. (2020). LncRNA Hmrhl regulates expression of cancer related genes in Chronic Myelogenous Leukemia through chromatin association. bioRxiv doi: https://doi.org/10.1101/2020.09.17.301770.

16. Cotney, J. L., and Noonan, J. P. (2015). Chromatin immunoprecipitation with fixed animal tissues and preparation for high-throughput sequencing. Cold Spring Harbor protocols, 2015(2), pdb–prot084848.

17. Cui, Y., Yin, Y., Xiao, Z., Zhao, Y., Chen, B., Yang, B., Xu, B., Song, H., Zou, Y., Ma, X. and Dai, J. (2019). LncRNA Neat1 mediates miR-124-induced activation of Wnt/β-catenin signaling in spinal cord neural progenitor cells. Stem cell research & therapy, 10(1), pp.1–11.

18. de Planell-Saguer, M., Rodicio, M. C., and Mourelatos, Z. (2010). Rapid in situ codetection of noncoding RNAs and proteins in cells and formalin-fixed paraffin-embedded tissue sections without protease treatment. Nature protocols, 5(6), 1061.

19. Derrien, T., Johnson, R., Bussotti, G., Tanzer, A., Djebali, S., Tilgner, H., Guernec, G., Martin, D., Merkel, A., Knowles, D.G. and Lagarde, J. (2012). The GENCODE v7 catalog of human long noncoding RNAs: analysis of their gene structure, evolution, and expression. Genome research, 22(9), pp.1775–1789.

20. Dinger, M.E., Amaral, P.P., Mercer, T.R., Pang, K.C., Bruce, S.J., Gardiner, B.B., Askarian-Amiri, M.E., Ru, K., Soldà, G., Simons, C. and Sunkin, S.M. (2008). Long noncoding RNAs in mouse embryonic stem cell pluripotency and differentiation. Genome research, 18(9), pp.1433–1445.

21. Engreitz, J.M., Haines, J.E., Perez, E.M., Munson, G., Chen, J., Kane, M., McDonel, P.E., Guttman, M. and Lander, E.S. (2016). Local regulation of gene expression by lncRNA promoters, transcription and splicing. Nature, 539(7629), pp.452–455.

22. Fatima, R., Choudhury, S.R., Divya, T.R., Bhaduri, U. and Rao, M.R.S. (2019). A novel enhancer RNA, Hmrhl, positively regulates its host gene, phkb, in chronic myelogenous leukemia. Non-coding RNA research, 4(3), pp.96–108.

23. Feng, J., Bi, C., Clark, B.S., Mady, R., Shah, P. and Kohtz, J.D. (2006). The Evf-2 noncoding RNA is transcribed from the Dlx-5/6 ultraconserved region and functions as a Dlx-2 transcriptional coactivator. Genes & development, 20(11), pp.1470–1484.

24. Fullard, J.F., Hauberg, M.E., Bendl, J., Egervari, G., Cirnaru, M.D., Reach, S.M., Motl, J., Ehrlich, M.E., Hurd, Y.L. and Roussos, P. (2018). An atlas of chromatin accessibility in the adult human brain. Genome research, 28(8), pp.1243–1252.

25. Gaiano, N., Nye, J. S., and Fishell, G. (2000). Radial glial identity is promoted by Notch1 signaling in the murine forebrain. Neuron, 26(2), 395–404.

26. Ganesan, G. and Rao, S.M. (2008). A novel noncoding RNA processed by Drosha is restricted to nucleus in mouse. Rna, 14(7), pp.1399–1410.

27. Goff, L.A., Groff, A.F., Sauvageau, M., Trayes-Gibson, Z., Sanchez-Gomez, D.B., Morse, M., Martin, R.D., Elcavage, L.E., Liapis, S.C., Gonzalez-Celeiro, M. and Plana, O. (2015). Spatiotemporal expression and transcriptional perturbations by long noncoding RNAs in the mouse brain. Proceedings of the National Academy of Sciences, 112(22), pp.6855–6862.

28. Grant, C. E., Bailey, T. L., and Noble, W. S. (2011). FIMO: scanning for occurrences of a given motif. Bioinformatics, 27(7), 1017–1018.

29. Hart, R.P. and Goff, L.A. (2016). Long noncoding RNAs: central to nervous system development. International Journal of Developmental Neuroscience, 55, pp.109–116.

30. Hartford, C.C.R. and Lal, A. (2020). When long noncoding becomes protein coding. Molecular and Cellular Biology, 40(6).

31. Hezroni, H., Ben-Tov Perry, R., Gil, N., Degani, N. and Ulitsky, I. (2020). Regulation of neuronal commitment in mouse embryonic stem cells by the Reno1/Bahcc1 locus. EMBO reports, 21(11), p.e51264.

32. Huang, T., Wang, G., Yang, L., Peng, B., Wen, Y., Ding, G. and Wang, Z. (2017). Transcription factor YY1 modulates lung cancer progression by activating lncRNA-PVT1. DNA and cell biology, 36(11), pp.947–958.

33. Imayoshi, I., Sakamoto, M., Yamaguchi, M., Mori, K., and Kageyama, R. (2010). Essential roles of Notch signaling in maintenance of neural stem cells in developing and adult brains. Journal of Neuroscience, 30(9), 3489–3498.

34. Iyer, M.K., Niknafs, Y.S., Malik, R., Singhal, U., Sahu, A., Hosono, Y., Barrette, T.R., Prensner, J.R., Evans, J.R., Zhao, S. and Poliakov, A. (2015). The landscape of long noncoding RNAs in the human transcriptome. Nature genetics, 47(3), pp.199–208.

35. Jarroux, J., Morillon, A. and Pinskaya, M. (2017). History, discovery, and classification of lncRNAs. Long Non Coding RNA Biology, pp.1–46.

36. Kadakkuzha, B.M., Liu, X.A., McCrate, J., Shankar, G., Rizzo, V., Afinogenova, A., Young, B., Fallahi, M., Carvalloza, A.C., Raveendra, B. and Puthanveettil, S.V. (2015). Transcriptome analyses of adult mouse brain reveal enrichment of lncRNAs in specific brain regions and neuronal populations. Frontiers in cellular neuroscience, 9, p.63.

37. Kamachi, Y., Uchikawa, M., Tanouchi, A., Sekido, R., and Kondoh, H. (2001). Pax6 and SOX2 form a co-DNA-binding partner complex that regulates initiation of lens development. Genes & development, 15(10), 1272–1286.

38. Kirkeby, A., Grealish, S., Wolf, D.A., Nelander, J., Wood, J., Lundblad, M., Lindvall, O. and Parmar, M. (2012). Generation of regionally specified neural progenitors and functional neurons from human embryonic stem cells under defined conditions. Cell reports, 1(6), 703–714.

39. Knauss, J.L., Miao, N., Kim, S.N., Nie, Y., Shi, Y., Wu, T., Pinto, H.B., Donohoe, M.E. and Sun, T. (2018). Long noncoding RNA Sox2ot and transcription factor YY1 co-regulate the differentiation of cortical neural progenitors by repressing Sox2. Cell death & disease, 9(8), pp.1–13.

40. Kopp, F. and Mendell, J.T. (2018). Functional classification and experimental dissection of long noncoding RNAs. Cell, 172(3), pp.393–407.

41. Lee, T. Y., Chang, W. C., Hsu, J. B. K., Chang, T. H., and Shien, D. M. (2012). GPMiner: an integrated system for mining combinatorial cis-regulatory elements in mammalian gene group. In BMC genomics, 13, 1, pp. 1–12.

42. Li, L., Zhuang, Y., Zhao, X. and Li, X. (2019). Long non-coding RNA in neuronal development and neurological disorders. Frontiers in genetics, 9, p.744.

43. Liau, W.S., Samaddar, S., Banerjee, S. and Bredy, T.W. (2021). On the functional relevance of spatiotemporally-specific patterns of experience-dependent long noncoding RNA expression in the brain. RNA biology, pp.1–12.

44. Liu, J., Wu, X., Zhang, H., Pfeifer, G. P., and Lu, Q. (2017). Dynamics of RNA polymerase II pausing and bivalent histone H3 methylation during neuronal differentiation in brain development. Cell reports, 20(6), 1307–1318.

45. Lv, J., Cui, W., Liu, H., He, H., Xiu, Y., Guo, J., Liu, H., Liu, Q., Zeng, T., Chen, Y. and Zhang, Y. (2013). Identification and characterization of long non-coding RNAs related to mouse embryonic brain development from available transcriptomic data. PloS one, 8(8), p.e71152.

46. Machon, O., Van Den Bout, C. J., Backman, M., Kemler, R., and Krauss, S. (2003). Role of β-catenin in the developing cortical and hippocampal neuroepithelium. Neuroscience, 122(1), 129–143.

47. Marchese, F.P., Raimondi, I. and Huarte, M. (2017). The multidimensional mechanisms of long noncoding RNA function. Genome biology, 18(1), pp.1–13.

48. Mathelier, A., Zhao, X., Zhang, A.W., Parcy, F., Worsley-Hunt, R., Arenillas, D.J., Buchman, S., Chen, C.Y., Chou, A., Ienasescu, H. and Lim, J. (2014). JASPAR 2014: an extensively expanded and updated open-access database of transcription factor binding profiles. Nucleic acids research, 42(D1), D142-D147

49. Mercer, T.R., Dinger, M.E., Sunkin, S.M., Mehler, M.F. and Mattick, J.S. (2008). Specific expression of long noncoding RNAs in the mouse brain. Proceedings of the National Academy of Sciences, 105(2), pp. 716-721.

50. Mizutani, K. I., Yoon, K., Dang, L., Tokunaga, A., and Gaiano, N. (2007). Differential Notch signalling distinguishes neural stem cells from intermediate progenitors. Nature, 449(7160), 351–355.

51. Mohamed, J.S., Gaughwin, P.M., Lim, B., Robson, P. and Lipovich, L. (2010). Conserved long noncoding RNAs transcriptionally regulated by Oct4 and Nanog modulate pluripotency in mouse embryonic stem cells. Rna, 16(2), pp.324–337.

52. Ng, S.Y., Bogu, G.K., Soh, B.S. and Stanton, L.W. (2013). The long noncoding RNA RMST interacts with SOX2 to regulate neurogenesis. Molecular cell, 51(3), pp.349–359.

53. Nishant, K.T., Ravishankar, H. and Rao, M.R.S. (2004). Characterization of a mouse recombination hot spot locus encoding a novel non-protein-coding RNA. Molecular and cellular biology, 24(12), p.5620.

54. Owa, T., Taya, S., Miyashita, S., Yamashita, M., Adachi, T., Yamada, K., Yokoyama, M., Aida, S., Nishioka, T., Inoue, Y.U. and Goitsuka, R. (2018). Meis1 coordinates cerebellar granule cell development by regulating Pax6 transcription, BMP signaling and Atoh1 degradation. Journal of Neuroscience, 38(5), 1277–1294.

55. Pal, D., Neha, C.V., Bhaduri, U., Zenia, Z., Dutta, S., Chidambaram, S. and Rao, M.R.S. (2021). LncRNA Mrhl orchestrates differentiation programs in mouse embryonic stem cells through chromatin mediated regulation. Stem Cell Research, 53, p.102250.

56. Pijnappel, W. P., Baltissen, M. P., and Timmers, H. M. (2013). Protocol for lentiviral knock down in mouse ES cells. Doi 10.1038/protex.2013.036

57. Pinson, J., Mason, J. O., Simpson, T. I., and Price, D. J. (2005). Regulation of the Pax6: Pax6 (5a) mRNA ratio in the developing mammalian brain. BMC developmental biology, 5(1), 1–4.

58. Plachta, N., Bibel, M., Tucker, K. L., and Barde, Y. A. (2004). Developmental potential of defined neural progenitors derived from mouse embryonic stem cells. Development, 131(21), 5449–5456.

59. Ramos, A.D., Andersen, R.E., Liu, S.J., Nowakowski, T.J., Hong, S.J., Gertz, C.C., Salinas, R.D., Zarabi, H., Kriegstein, A.R. and Lim, D.A. (2015). The long noncoding RNA Pnky regulates neuronal differentiation of embryonic and postnatal neural stem cells. Cell stem cell, 16(4), pp.439–447.

60. Roberts, T.C., Morris, K.V. and Wood, M.J. (2014). The role of long non-coding RNAs in neurodevelopment, brain function and neurological disease. Philosophical Transactions of the Royal Society B: Biological Sciences, 369(1652), p.20130507.

61. Rosa, A. and Ballarino, M. (2016). Long noncoding RNA regulation of pluripotency. Stem Cells International, 2016.

62. Sakamoto, M., Hirata, H., Ohtsuka, T., Bessho, Y., and Kageyama, R. (2003). The basic helix-loop-helix genes Hesr1/Hey1 and Hesr2/Hey2 regulate maintenance of neural precursor cells in the brain. Journal of Biological Chemistry, 278(45), 44808–44815.

63. Sansom, S.N., Griffiths, D.S., Faedo, A., Kleinjan, D.J., Ruan, Y., Smith, J., Van Heyningen, V., Rubenstein, J.L. and Livesey, F.J. (2009). The level of the transcription factor Pax6 is essential for controlling the balance between neural stem cell self-renewal and neurogenesis. PLoS Genet, 5(6), e1000511.

64. Stamou, M., Ng, S.Y., Brand, H., Wang, H., Plummer, L., Best, L., Havlicek, S., Hibberd, M., Khor, C.C., Gusella, J. and Balasubramanian, R. (2020). A balanced translocation in Kallmann Syndrome implicates a long noncoding RNA, RMST, as a GnRH neuronal regulator. The Journal of Clinical Endocrinology & Metabolism, 105(3), pp.e231–e244.

65. Su, Z., Zhang, Y., Liao, B., Zhong, X., Chen, X., Wang, H., Guo, Y., Shan, Y., Wang, L. and Pan, G. (2018). Antagonism between the transcription factors NANOG and OTX2 specifies rostral or caudal cell fate during neural patterning transition. Journal of Biological Chemistry, 293(12), 4445–4455.

66. Sun, J., Rockowitz, S., Xie, Q., Ashery-Padan, R., Zheng, D., and Cvekl, A. (2015). Identification of in vivo DNA-binding mechanisms of Pax6 and reconstruction of Pax6-dependent gene regulatory networks during forebrain and lens development. Nucleic acids research, 43(14), 6827–6846.

67. Sun, J., Zhao, Y., McGreal, R., Cohen-Tayar, Y., Rockowitz, S., Wilczek, C., Ashery-Padan, R., Shechter, D., Zheng, D. and Cvekl, A. (2016). Pax6 associates with H3K4-specific histone methyltransferases Mll1, Mll2, and Set1a and regulates H3K4 methylation at promoters and enhancers. Epigenetics & chromatin, 9(1), pp.1–18.

68. Sun, M. and Kraus, W.L. (2015). From discovery to function: the expanding roles of long noncoding RNAs in physiology and disease. Endocrine reviews, 36(1), pp.25–64.

69. Sun, Z., Huang, G. and Cheng, H. (2019). Transcription factor Nrf2 induces the upregulation of lncRNA TUG1 to promote progression and adriamycin resistance in urothelial carcinoma of the bladder. Cancer management and research, 11, p.6079.

70. Thakurela, S., Tiwari, N., Schick, S., Garding, A., Ivanek, R., Berninger, B., and Tiwari, V. K. (2016). Mapping gene regulatory circuitry of Pax6 during neurogenesis. Cell discovery, 2(1), 1–22.

71. Tian, K., Wang, A., Wang, J., Li, W., Shen, W., Li, Y., Luo, Z., Liu, Y. and Zhou, Y. (2021). Transcriptome Analysis Identifies SenZfp536, a Sense LncRNA that Suppresses Self-renewal of Cortical Neural Progenitors. Neuroscience Bulletin, 37(2), pp.183–200.

72. Uchil, P. D., Nagarajan, A., and Kumar, P. (2017). Assay for β-galactosidase in extracts of mammalian cells. Cold Spring Harbor Protocols, 2017(10), pdb–prot095778.

73. Vihma, H., Luhakooder, M., Pruunsild, P., and Timmusk, T. (2016). Regulation of different human NFAT isoforms by neuronal activity. Journal of neurochemistry, 137(3), 394–408.

74. Wang, H., Huo, X., Yang, X.R., He, J., Cheng, L., Wang, N., Deng, X., Jin, H., Wang, N., Wang, C. and Zhao, F. (2017). STAT3-mediated upregulation of lncRNA HOXD-AS1 as a ceRNA facilitates liver cancer metastasis by regulating SOX4. Molecular cancer, 16(1), 1–15.

75. Wapinski, O. and Chang, H.Y. (2011). Long noncoding RNAs and human disease. Trends in cell biology, 21(6), pp.354–361.

76. Wild, A. R., Sinnen, B. L., Dittmer, P. J., Kennedy, M. J., Sather, W. A., and Dell’Acqua, M. L. (2019). Synapse-to-Nucleus communication through NFAT is mediated by L-type Ca2+ Channel Ca2+ Spike Propagation to the Soma. Cell reports, 26(13), 3537–3550.

77. Wilusz, J.E., Sunwoo, H. and Spector, D.L. (2009). Long noncoding RNAs: functional surprises from the RNA world. Genes & development, 23(13), pp.1494–1504.

78. Woodhead, G. J., Mutch, C. A., Olson, E. C., and Chenn, A. (2006). Cell-autonomous β-catenin signaling regulates cortical precursor proliferation. Journal of Neuroscience, 26(48), 12620–12630.

79. Wu, S.C., Kallin, E.M. and Zhang, Y. (2010). Role of H3K27 methylation in the regulation of lncRNA expression. Cell research, 20(10), pp. 1109–1116.

80. Wu, Z., Liu, X., Liu, L., Deng, H., Zhang, J., Xu, Q., Cen, B. and Ji, A. (2014). Regulation of lncRNA expression. Cellular & molecular biology letters, 19(4), pp.561–575.

81. Xie, Q., and Cvekl, A. (2011). The orchestration of mammalian tissue morphogenesis through a series of coherent feed-forward loops. Journal of Biological Chemistry, 286(50), 43259–43271.

82. Xie, Q., Yang, Y., Huang, J., Ninkovic, J., Walcher, T., Wolf, L., Vitenzon, A., Zheng, D., Götz, M., Beebe, D.C. and Zavadil, J. (2013). Pax6 interactions with chromatin and identification of its novel direct target genes in lens and forebrain. PLoS One, 8(1), e54507.

83. Xing, J., Liu, H., Jiang, W. and Wang, L. (2020). LncRNA-encoded peptide: functions and predicting methods. Frontiers in Oncology, 10.

84. Xu, Y., Xi, J., Wang, G., Guo, Z., Sun, Q., Lu, C., Ma, L., Wu, Y., Jia, W., Zhu, S. and Guo, X. (2021). PAUPAR and PAX6 sequentially regulate human embryonic stem cell cortical differentiation. Nucleic acids research, 49(4), pp.1935–1950.

85. Yamada, T., Urano-Tashiro, Y., Tanaka, S., Akiyama, H., and Tashiro, F. (2013). Involvement of crosstalk between Oct4 and Meis1a in neural cell fate decision. PLoS One, 8(2), e56997.

86. Yao, B., and Jin, P. (2014). Unlocking epigenetic codes in neurogenesis. Genes & development, 28(12), 1253–1271.

87. Yoon, J.H., Abdelmohsen, K. and Gorospe, M. (2013). Posttranscriptional gene regulation by long noncoding RNA. Journal of molecular biology, 425(19), pp.3723–3730.

88. Zelinger, L., Karakülah, G., Chaitankar, V., Kim, J.W., Yang, H.J., Brooks, M.J. and Swaroop, A. (2017). Regulation of noncoding transcriptome in developing photoreceptors by rod differentiation factor NRL. Investigative ophthalmology & visual science, 58(11), pp.4422–4435.

89. Zhang, P., Wu, W., Chen, Q. and Chen, M. (2019, a). Non-coding RNAs and their integrated networks. Journal of integrative bioinformatics, 16(3).

90. Zhang, L., Xue, Z., Yan, J., Wang, J., Liu, Q. and Jiang, H. (2019, b). LncRNA Riken-201 and Riken-203 modulates neural development by regulating the Sox6 through sequestering miRNAs. Cell proliferation, 52(3), p.e12573.

91. Zhang, L., Yan, J., Liu, Q., Xie, Z. and Jiang, H. (2019, c). LncRNA Rik-203 contributes to anesthesia neurotoxicity via microRNA-101a-3p and GSK-3β-mediated neural differentiation. Scientific reports, 9(1), pp.1–12.

92. Zhao, Y., Liu, H., Zhang, Q. and Zhang, Y. (2020). The functions of long non-coding RNAs in neural stem cell proliferation and differentiation. Cell & Bioscience, 10, pp.1–10.

